# Predicting continuous outcomes: Some new tests of associative approaches to contingency learning

**DOI:** 10.1101/2025.06.12.659290

**Authors:** Julie Y. L. Chow, Hilary J. Don, Ben Colagiuri, Evan J. Livesey

## Abstract

Associative learning models have traditionally simplified contingency learning by relying on binary classification of cues and outcomes, such as administering a medical treatment (or not) and observing whether the patient recovered (or not). While successful in capturing fundamental learning phenomena across human and animal studies, these models are not capable of representing variability in human experiences that are common in many real-world contexts. Indeed, where variation in outcome magnitude exists (e.g., severity of illness in a medical scenario), this class of models, at best, approximate the outcome mean with no ability to represent the underlying *distribution* of values. In this paper, we introduce one approach to incorporating a distributed architecture into a prediction error learning model that tracks the contingency between cues and dimensional outcomes. Our Distributed Model allows associative links to form between the cue and outcome nodes that provide distributed representation depending on the magnitude of the outcome, thus enabling learning that extends beyond approximating the mean. Comparing the Distributed Model against a Simple Delta Model across four contingency learning experiments, we found that the Distributed Model provides significantly better fit to empirical data in virtually all participants. These findings suggest human learners rely on a means of encoding outcomes that preserves the continuous nature of experienced events, advancing our understanding of causal inference in complex environments.

**Author Summary:** When we learn about cause and effect in everyday life—such as whether a medicine helps recovery from illness—we experience outcomes that vary in degree rather than simply happening or not happening. Traditional models of how humans and animals learn have largely focused on these all-or-nothing scenarios, essentially tracking the average value when outcomes are dimensional. We developed a model that extends on simple error-correction models to represent how people learn about relationships between cues and outcomes that can take on a range of values. Instead of just tracking the average, our Distributed Model captures the full spectrum of possible outcomes and their frequencies. We tested this model against a conventional single point-estimate approach across four experiments and found that our Distributed Model better matched how people make predictions in nearly every case. Our findings suggest that a relatively simple adjustment to conventional prediction-error learning algorithms that allows representation of outcome magnitudes provide a powerful way to capture the information that we preserve when we learn about variable outcomes. This has important implications for understanding how people make predictions and decisions in real-world situations where outcomes naturally vary, from medical treatments to environmental changes.

## Introduction

Contemporary models of associative learning are founded on their ability to reproduce phenomena observed across the animal kingdom and, critically, this includes effects that translate well to human contingency learning. Over the last 50 years, a wealth of contingency learning data has accumulated, tackling issues such as the sensitivity of causal judgments to contingency (Jenkins & Ward, 1965), biases in perceiving contingent relationships when none exist (e.g., Hamilton & Gifford, 1976; Matute et al., 2015), and cue competition (e.g., Kamin, 1969). The vast majority of these data come from experiments with simple and discrete stimuli that do not vary substantially in magnitude. Outcomes are usually visually represented as being present or absent, enabling the degree of association between a cue and an outcome to be determined by the covariation of binary events. This is, of course, a dramatic simplification of the events that a learner encounters in the real world, which often vary by degree rather than merely being present or absent. To illustrate, contingency learning tasks using a medical scenario typically involve presenting a medical treatment and evaluating whether the patient has recovered or not recovered (i.e., two possible outcome states; e.g., Vinas et al., 2025). Most health experiences however, involve fluctuating and noisy information and it is often unclear how one might characterise these experiences in a dichotomous way. The use of binary outcomes is convenient and justifiable in many circumstances (e.g., it helps to ensure we have theoretically tractable research questions), but it potentially misses a rich source of variation in the way we learn and avoids addressing challenges that models of learning must face in order to provide an adequate account of how people infer causal relationships in the real world. While contingency learning tasks may not always require participants to make explicit causal judgments—that is, whether the cue *causes* the outcome to occur—the process of learning predictive relationships from contingencies forms an important foundation for causal inference (see Shanks & Dickinson, 1988).

More recently, much has been made about learning complex properties, including distributional learning (people find it difficult to reproduce experienced distributions, e.g., Tran et al., 2017), and uncertainty related to making predictions (e.g., Courville et al., 2006; Dayan et al., 2000). Research examining how people learn from continuous and variable outcomes has developed along multiple trajectories, with different modelling approaches emerging to address distinct theoretical questions and task demands. A Bayesian approach to causal inference explicitly represents uncertainty in its model, and this approach has been extended from its original remit, predicting binary variables (Griffiths & Tenenbaum, 2005), to continuous ones (Pacer & Griffiths, 2011). However, for the most part, the Bayesian approach is concerned with defining and formalising the problem of causal inference, that is, *what* should be calculated for valid causal induction. These computational-level models provide normative accounts of how ideal observers should reason about causal structure, but they do not necessarily specify the mechanisms by which humans actually perform this inference. At the other extreme, mechanistic models, including the prediction error learning networks we use in this study, are concerned with providing some account of the psychological mechanisms responsible for mental representation of events and how we learn about them. Some approaches bridge these levels by implementing Bayesian principles with prediction error mechanisms, such as the Hierarchical Gaussian Filter (Mathys et al., 2011, 2014; Mikus et al., 2025), which uses prediction errors to update hierarchical beliefs about hidden environmental states. While mechanistic in their implementation, such models still assume substantial computational sophistication on the part of the learner. For instance, learners must track not only their current beliefs about outcomes but also their confidence in those beliefs, distinguish between stable noise and genuine environmental changes, and adjust how much they learn based on these uncertainty estimates across multiple hierarchical levels simultaneously. Mechanistic models built on Bayesian principles typically assume that the learner represents uncertainty as a distinct variable and performs computational operations on it, which goes substantially beyond simple error-driven learning. In contrast, we explore whether learning about continuous outcomes – and the variability in responding that arises from these experiences – can emerge from simple associative mechanisms without any explicit uncertainty representation or computation.

### Associative approach to contingency learning

Connectionist models assume that people track the covariation between events through mental links connecting representations of these events. The strength of these connections change with increasing experience. Many use a simple error-correction learning algorithm like that proposed by Rescorla and Wagner (1972; the RW model) in which learning is proportional to the discrepancy between what is expected and what actually occurs (i.e., *prediction error*). With increased experience of a putative cause (commonly referred to as a cue) and outcome co-occurring, associative weights linking the two events strengthen, such that the outcome is increasingly expected when the cue is present. Predictions about the outcome can be directly estimated by taking the summed associative strength of all cues present (i.e., how much the outcome is expected given cues present). Violation of expectancy—for example, when the outcome is expected but does not occur—produces a prediction error signal that leads to updating of associative strengths, i.e., learning. Thus, on each trial the model provides an estimate of the associative strength, and this estimate is updated based on the whether the outcome is present or absent.

The appeal of simple connectionist models like the RW model is that the process of learning is greatly simplified by providing a single point estimate for the outcome. In a contingency learning task where the outcome has some numeric value that changes trial-to-trial (e.g., magnitude of symptom severity), the associative strength will approximate the outcome mean. The implication of this is that prediction error models of this class are models of the outcome mean; they are not typically designed to provide an accurate fit of the *distribution* of outcomes, where variation in outcome magnitude exists, thereby missing a rich source of information concerning how we learn about variable events. However, there is scope for these models to represent outcomes in a distributed network. The present paper demonstrates one way in which a distributed representation of outcome values can be incorporated into a prediction error model of contingency learning.

### Distributional representation in learning

Distributed stimulus representation is not new in associative learning. However, it is most commonly used to represent predictive *cues* that vary along a continuum (e.g., McLaren & Mackintosh, 2002; Livesey & McLaren, 2019; Thorwart, Harris & Livesey, 2012; Ghirlanda & Enquist, 1998). Relatively few studies within the human associative learning tradition have examined how well people learn about continuous *outcomes*. Previous work from our laboratory has shown that some phenomena typically found in contingency learning research using discrete outcomes were also present when participants were presented with continuous outcomes (e.g., outcome density effect; Chow et al., 2019; Double et al., 2020), suggesting that a common cognitive process may be involved in learning about discrete and dimensional outcomes. In some instances, researchers have used continuous outcomes to leverage a certain type of cognitive process. For instance, some of the debates about whether people show blocking for reasons to do with deductive reasoning or other decision processes make use of outcomes that are demonstrably additive and/or submaximal on some magnitude scale (Beckers et al., 2005; Lovibond et al., 2003; Livesey & Boakes, 2004; Livesey et al., 2019). In these instances, however, variations in magnitude are a means to an end; these experiments are not concerned with whether people learn about continuous outcomes *per se*.

In contrast, substantive work has been done using predictive inference tasks, examining how people track outcomes that vary across trials, often in environments where the statistics of those outcomes change over time (e.g., Nassar et al., 2010, 2012, 2021; Piray & Daw, 2021, 2024). These paradigms differ from contingency learning tasks in that the primary goal is to detect when shifts in outcome statistics have occurred (often referred to as environmental volatility) and adjust predictions accordingly. Models developed from these tasks have demonstrated that simpler mechanistic alternatives to full Bayesian inference (Nassar et al., 2010) and models with adaptive learning rates (Piray & Daw, 2021, 2024), can capture key patterns of human behaviour without the full computational machinery of hierarchical Bayesian inference. Critically, these models typically represent predictions as single point estimates (a current best guess about the outcome mean) with learning rates that adapt based on various uncertainty signals. While this approach is well-suited for tasks requiring rapid detection of environmental changes, it may be less appropriate for contingency learning contexts where the goal is not to track a changing mean but to learn about stable cue-outcome relationships from accumulated experiences. These different task demands raise the question of whether models that preserve the distributional nature of experienced outcomes better approximate the way people represent and learn about continuous outcomes in contingency learning tasks. For instance, people might draw from a range of experienced outcomes when evaluating whether a treatment is effective, not just their most recent experience. Understanding how learners represent and utilise distributional information in contingency learning remains an open question, and one that may require different modelling approaches than those developed for predictive inference tasks.

Distributed representations have previously been explored in other learning domains. For example, in function learning, the learner is asked to predict the magnitude of the outcome based on the level of input. The functional relationship between cue and outcome can be determined by some formula (e.g., a linear relationship) plus some noise (DeLosh et al., 1997; McDaniel et al., 2014): as the input at the cue level increases, the magnitude of the output also increases to a similar degree. Function learning has a place in everyday life, for example a doctor who is trying to estimate the change in the patient’s body temperature given the severity of infection. This type of learning allows people to understand how the magnitude of an effect changes according to the strength of the cause, and thus make reliable predictions based on that relationship. There is evidence that people readily learn these functional relationships (Byun, 1995), and that knowledge can be captured in an associative framework (Kruschke, 1992; DeLosh et al., 1997). Associative learning models in function learning provide a starting point to consider how a distributed network for representing complex properties of outcomes can be implemented in contingency learning models.

The aim of the current project is to develop a connectionist model of contingency learning capable of representing continuous outcomes in a distributed fashion, and formalise a way in which distributions of responses might be derived from such a model. In particular, we are interested in what can be gained from a distributed approach to representing continuous outcomes in a prediction-error learning model, compared to a similar model which produces only a single point estimation that approximates the outcome mean. We will first discuss one approach that could be used to represent distribution of outcome values, and then report the outcomes of a formal model comparison comparing the distributed model against a simple connectionist model that does not implement distributed outcome representation.

## Methods

### Modelling continuous outcomes with a prediction error algorithm

#### Simple Delta Model

We present here a description of a simple prediction error model (or delta rule model) commonly used to model trial-by-trial learning. The basic feature of a delta rule model is that the strength of connection between a single cue and outcome is updated on each trial according to a learning algorithm like the one shown in Equation 1:

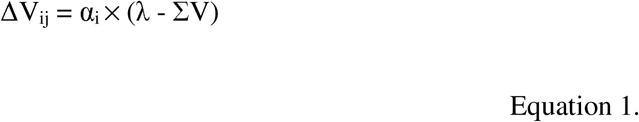

In this equation, the change in the strength of the connection between cue i and the outcome j, changes according to prediction error, that is, the discrepancy between the observed outcome, λ, and the extent to which it is predicted by the cues present, ΣV. Note that among the cue representations, it is assumed that the context is also represented by one or several nodes in the connectionist network, entering into associations with the outcome like other cue nodes. In all the simulations we present here, we used a single cue node for each predictive cue plus one additional cue node for the context. Since the four experiments presented in this study all used single-cue trials during learning, this means that ΣV was always the sum of the association strength of the presented cue node plus the association strength of the context node (this also holds true for calculating the individual ΣV for each outcome node in the distributed model).

When cues are followed by the outcome, (λ – ΣV) is typically positive, though the amount that will be learned about their co-occurrence depends on whether the outcome is surprising (larger prediction error = more learning) or expected (smaller prediction error = less learning). When cues are followed by the *omission* of the outcome, λ - ΣV can be negative if the outcome was expected. That is, if ΣV is greater than λ, then the connection linking the cue and outcome is weakened, or becomes increasingly inhibitory if V < 0. This simple interaction between expectation and error allows the algorithm to track contingencies between cues and outcomes accurately. Additional parameter α denotes the learning rate of each node with the outcome: α for each individual cue node and α_x_ for the context node, which is given an independent value because its properties are, by definition, different to the cues. The simplest instantiations of this model, those used to model a single outcome with two states (present vs absent) for instance, assume that the absence of the outcome does not have its own unique representation – it is simply represented as a situation in which λ = 0 in Equation 1.

In the present paper, as a starting point we assume that the representation of the outcome itself must have some relationship with the outcomes that are observed on each trial. Thus, on each trial λ takes on the observed outcome value, which in our case is a value between 0 and 100. We make the additional assumption here that participants are biased towards the center of the outcome scale at the start of the study, thus the associative strength V for the context starts at a value of 50. In all other respects, the model operates like a conventional delta rule model, and V for the cue approximates the mean outcome value at that point in time. Both associative strengths of the cue and the context update independently on trials where the cue is present, but only the associative strength of the context is updated when the cue is absent.

Delta rule models usually assume that the prediction on each trial is determined by ΣV (or at least a monotonic function of ΣV). However, our preliminary investigations suggested that relying on ΣV alone produced unrealistically low variance in predictions; for a particular trial type, ΣV itself varies to a degree across trials depending on the outcomes experienced, but the prediction will always reflect ΣV on that trial. We thus additionally introduced a noise parameter, *σ_e_*to capture unsystematic variability in *predictions* based on the learner’s estimate of expected value. In effect, this means that the learner had some chance of choosing values that were similar to the expected value ΣV, even though learning tracked a single value. To achieve this, we assumed that a prediction of value *r* had a choice weight *W* created using a gaussian function with a probability distribution centered around the expected value ΣV, with a variance of *σ_e_^2^*.

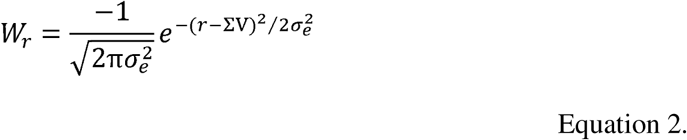

Unlike the Distributed Model which we will present below, the variability introduced with the *σ_e_* is not implemented at the learning stage but only used to generate simulated predictions. Note that we use the notation *σ_e_* to denote the variation in *explicit predictions* added to the Simple Delta Model, and *σ_n_* to denote the breadth of *node* activation in the outcome representations in the Distributed Model. The theoretical assumption in the Simple Delta Model is that participants are solely tracking the outcome mean during learning, but may still produce predictions that vary around that mean. In contrast, a distributed model would encode that variability *during learning*, resulting in a distribution of activation across outcome nodes. A corollary of this is that when generating simulated predictions with the Simple Delta Model, response probabilities are always normally distributed around ΣV, whereas in the Distributed Model response probabilities are, in principle, capable of demonstrating sensitivity to the shape and skew of the underlying outcome distribution.

In generating simulated predictions, we first assume that, all things being equal, the outcome value the person observes will be reflected in the predictions they subsequently make. Thus, like the participants in our experiments, the model can predict any integer value from 0 to 100 (101 distinguishable responses). In modelling human choices, the problem that is commonly faced is how discrete predictions are made on each trial. In many contexts, especially those involving discrete choices among several options, researchers assume that predictions are probabilistically determined. The *softmax* rule is often used for this purpose (Bridle, 1990; Rumelhart et al., 1995):

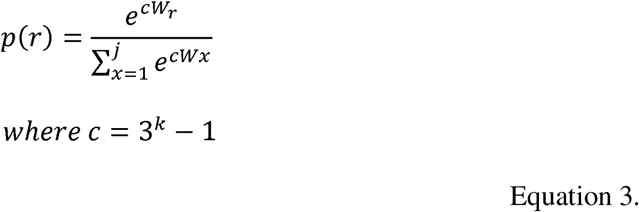

Equation 3 calculates the probability of choosing a response, *r* from a set of j choices (the probabilities of all choices summing to 1). The probability of each choice, *p*(*r*) is determined by the choice weight *W*_r_ as defined in Equation 2, and by an inverse temperature parameter *k*, which is scaled nonlinearly as *c* = 3*^k^*-1, following Don et al. (2019). This exponential transformation allows *c* to span a wide range of values while maintaining a bounded search space for *k* (described below), effectively placing an implicit prior on the inverse temperature with a truncated exponential distribution. The argument to the exponential function in the *softmax* calculation was additionally capped at 700, beyond which standard computational environments including R produce undefined values.

Values of *k* are restricted to a range from 0.001 to 5 (yielding positive values of *c* up to a maximum of 242) as is typical in the choice literature (e.g., Don et al., 2019). These constraints were implemented by specifying lower and upper bounds in the maximum likelihood estimation routine (*L-BFGS-B* in R’s `optim` function; see Byrd et al., 1995), such that the optimiser searched within this parameter space. If the gradient search points outside of the parameter space, the algorithm projects the step back to the boundary. The parameter *k* determines the stochasticity in choice behaviour: higher values of *k* lead to more consistent choice of the outcome node with the highest activation, whereas when *k* is a small value, choice probabilities become increasingly similar and a large difference in relative activation is required to produce a choice response with greater uncertainty (Yechiam & Ert, 2007). In short, the greater the relative magnitude of outcome node activation, the greater the probability of choosing the response. Using this algorithm, the Simple Delta Model is able to produce distribution-like responses despite learning only a point estimate. In addition, the approach is psychologically plausible: If participants were simply tracking the outcome mean with some uncertainty about that estimate, we would expect their responses to cluster around the learned mean with some noise.

We note briefly here that a feature of *softmax* is that it treats outcome nodes as independent, discrete choices, which may be a simplification given that participants are responding on a continuous scale (where nearby values may be more similar than values further apart). We acknowledge alternative modelling choices exist, including the use of a continuous response model, however we argue that these approaches cannot be straightforwardly applied to the Distributed Model, which produces a pattern of activations across nodes rather than a single point estimate with variance that could feed into a standard continuous response model. Forcing the Distributed Model into producing point estimates would also defeat the purpose of constructing a model capable of maintaining some fidelity to the distributed nature of the experienced outcomes. Thus, on the whole, we believe that the current approach provides a fair basis for comparing the two models. The *softmax* rule allows both models to generate discrete trial-by-trial predictions while preserving their theoretical differences, and gives the Simple Delta Model sufficient flexibility to produce variable responses comparable to those generated by the Distributed Model.

Finally, we include α for the cue and the context to be separate parameters determined by model fit, where α is the learning rate of the cue, and α_x_ is the learning rate of the context. In some instantiations of such a model, learning rate of the context, α_x_ is assumed to always be less than the learning rate of the cue, α; this constraint was not applied to model fit in the current study.

### Distributed Model

In this section, we will introduce one way in which distributed outcome representation can be incorporated into a delta rule model. One important difference between a simple delta rule approach and a distributed network approach is that in the former, there is a single associative value representing the strength of the connection between the cue and the outcome. In a distributed approach on the other hand, the cue forms an associative link with *multiple* outcome nodes along the dimension. Although it is not strictly necessary, for convenience, we used the same number of nodes as there are possible outcome values in our experiment. Thus, in the Distributed Model, there are as many associative links for each cue as there are outcome values. Learning then changes the strength of the connections between the cue and the outcome nodes. The strength of each connection depends on both the mean outcome value aggregated across trials, and the underlying distribution. This method of representing the outcome in a distributed network effectively tracks the occurrence of outcomes in a way that can differentiate between different outcome distributions that possess the same mean.

We incorporate a distributed architecture into the delta model by assuming that any given outcome magnitude is represented in a distributed fashion, resulting in activation of a series of outcome nodes that have overlapping sensitivities (see Figure 1). Activation of each node is a function of the distance between the observed outcome on that trial, and the peak sensitivity of the node. Each node is maximally sensitive to a particular outcome value, and activated incrementally less for outcomes that are larger or smaller than this value. For example, when the observed outcome has a value of 50, the outcome node corresponding to 50 is maximally activated. Values around 50 may also receive some level of activation, however node activation decreases as we move further away from the outcome node corresponding to the observed value. On every trial when an outcome value is presented, activity *a* in each outcome node *j* is determined according to a gaussian function, with a mean equal to the outcome value on that trial, λ, and variance around the mean, *σ_n_^2^*.

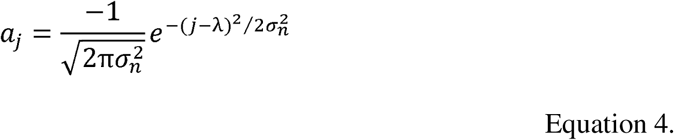

**Figure 1.**
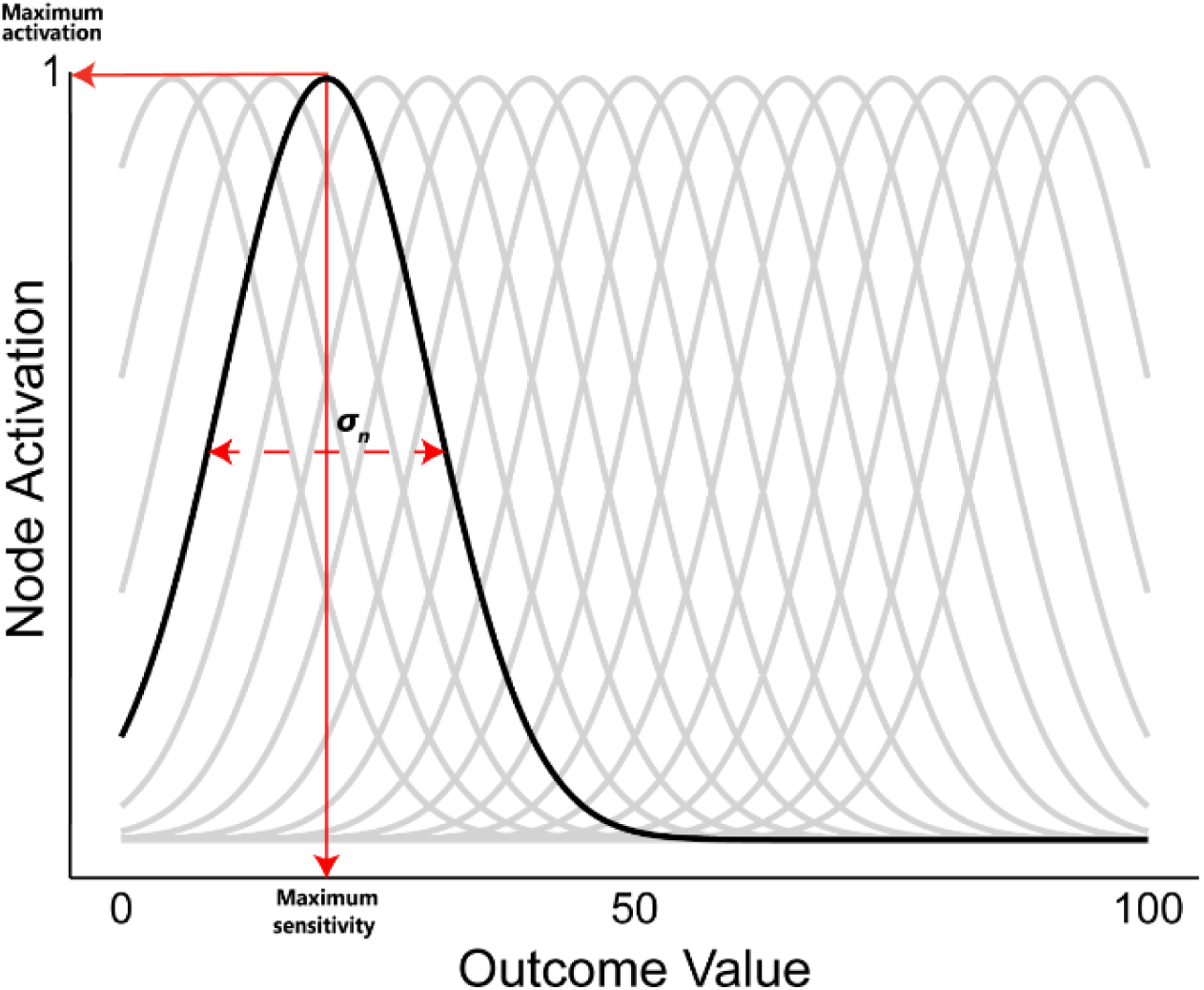
Illustration of distributed architecture in the Distributed Model, where the presentation of an outcome value will result in maximal activation at the outcome node corresponding to that value with decreasing activation away from that node following a gaussian function with variance, σ_n_. Fixed parameters are presented as a solid line, and estimated parameters as a dashed line.

The activation of these units is then normalised across all n outcome nodes so that they sum to 1, producing activation A_j_.

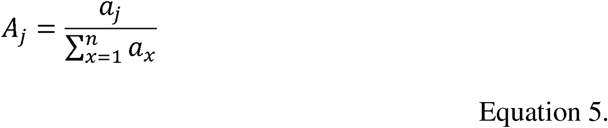

A corollary of this distributed architecture is that, unlike the Simple Delta Model (where we added unsystematic variance determined by *σ_e_*), choice weights for making predictions can be directly determined based on the ΣV of the corresponding outcome node (i.e., *W*_x_ = ΣV_x_) and thus used in the *softmax* rule as per Equation 3.

Associative learning for connections between each cue node and each outcome node then proceeds in the same way as for the simple delta rule, with the exception that the teaching signal for the learning algorithm is the normalised activation A_j_ produced by the observed outcome rather than by the observed value λ itself.

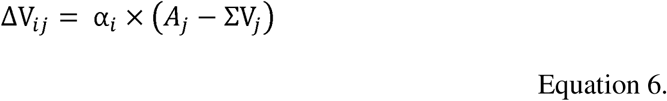

In contrast to the Simple Delta Model where the starting V for the context was set to outcome value of 50, starting values for V across the outcome dimension in the Distributed Model was randomly determined from values ranging between -.001 and +.001. Note that this difference in starting point should produce a greater advantage for the Simple Delta Model since participants tend to provide ratings at the mid-point of the scale. As we will see in the following section, the added advantage did not produce significantly better model fit for the Simple Delta Model. The strength of each associative link is updated on each trial following a prediction error algorithm in a similar fashion to the Simple Delta Model, that is by taking the difference in the outcome node activation and the expected value.

We fit each model to individual participants’ predictions made across the experiment using Maximum Likelihood Estimation (MLE). MLE selects parameter values that maximise the likelihood that the model produced the data observed in the empirical study. Both the Simple Delta and Distributed Model consisted of four estimated parameters required for model fitting and/or generating model predictions: context alpha (α_x_), cue alpha (α), variance in a gaussian function for making predictions in the Simple Delta Model (*σ_e_*) or activation spread during learning (*σ*_n_ in the Distributed Model), and temperature parameter for softmax, *k* required for model predictions. Best fitting parameters for both models for each participant in all four experiments can be found on the Open Science Framework (Chow et al., 2025).

In summary, both the Simple Delta and Distributed Model consist of the same number of estimated parameters for model fitting and making simulated predictions. Both models similarly rely on a prediction-error algorithm to update learning on each trial. The primary distinction between the two models is the ability for the Distributed Model to produce an associative link between the cue and each outcome node in the dimension (e.g., every value from 0 to 100 on a continuous scale). The relative activation of each outcome node across the outcome dimension allows the model to capture the underlying distribution of outcome values. In contrast, the Simple Delta Model updates a single associative value on every trial, producing a single terminal value (after many samples) that approximates the outcome mean; this approach has no ability to distinguish between cues with the same outcome mean but different underlying distributions (e.g., normal distribution vs negatively-skewed distribution).

In the following section, we will briefly describe a series of experiments and the empirical data used to determine model fit. We fitted both models using the same trial sequence and outcomes presented to participants in Experiments 1-4. We will provide a comparison of the two models across all four experiments using the Bayesian Information Criterion (BIC; Schwarz, 1978), followed by a discussion on model performance and simulated predictions.

## Empirical Data

We report here four empirical studies designed to determine participants’ sensitivity to outcome distributions. We designed these studies to include various factors that might influence learning, such as the presence of variability (Experiment 1), overall base-rate of outcome values (Experiment 2), and shape of the distribution in which outcome values are sampled from (Experiments 3 & 4). The purpose of these experiments was to obtain participant data that will later be used for model fit: we are interested in comparing the fit of both models (using best fitting parameters for each individual participant) to determine whether participants’ prediction ratings during learning were better captured by the Simple Delta Model or the Distributed Model that is capable of representing the underlying distribution of outcome values. For brevity, we report only an overview of the design of each experiment and findings here; detailed description of the study design (pp 2-6) and empirical data can be found in Supplementary Materials (pp 6-16). All procedures reported here were approved by The University of Sydney Human Research Ethics Committee (Approval Number 2015/670). All participants provided written consent to participate.

### Design & Procedure

Participants in all four experiments underwent a contingency learning experiment, where they were told to imagine that they were a medical researcher investigating a new illness and the efficacy of several fictitious drugs at treating this disease. Participants were shown a series of patients given one of the fictitious drug treatments and their subsequent health outcome on a scale from 0 (no recovery) to 100 (full recovery). The number of fictitious drug treatments differed between experiments (minimum = 8 [Experiment 3], maximum = 13 [Experiment 2; including filler cues]). Target cues were organised by distribution and mean outcome value. The allocation of stimulus name and image to cue type was random across participants. See Table 1 for a summary of the design of each experiment.

**Table 1.** Summary of the design of each experiment. Number in brackets represent the number of levels in the corresponding within-subjects factor.

| Experiment | Design |  |  |
| --- | --- | --- | --- |
|  | Target cues | Non-target cues | Total number of prediction trials |
| 1 | Mean (3): 25, 50, 75 x<br>Distribution (3): Fixed values, Uniform, Normal | Fixed outcome 0 and<br>Fixed outcome 100 | 11 cues x 21 repeats = 231 trials |
| 2 | Mean (5): 15, 30, 50, 75, 80<br>All cues had outcomes sampled from a normal distribution | Six filler cues for between-subjects base-rate manipulation: 3 x Base-rate condition 30, 50, 70<br>Fixed outcome 0 and<br>Fixed outcome 100 | ([5 target cues + 2 fixed-outcome cues] x 25 repeats) + (6 filler cues x 50 repeats) = 475 trials |
| 3 | Mean (2): 35, 65 x<br>Distribution (3): Normal, Positive-skew, Negative-skew | Fixed outcome 0 and<br>Fixed outcome 100 | (6 target cues x 40 repeats) + (2 fixed-outcome cues x 20 repeats) = 280 trials |
| 4 | Mean (2): 35, 65 x<br>Distribution (3): Bimodal, Normal- Narrow (Low Variance), Normal-Wide (High Variance) | Fixed outcome 0 and<br>Fixed outcome 100 | (6 target cues x 40 repeats) + (2 fixed-outcome cues x 20 repeats) = 280 trials |

#### General procedure of all experiments

Schematic of a single trial sequence in the learning phase is presented in Figure 2. Participants were told that each trial represented a different patient who was given one of the drug treatments. Participants were first shown the drug treatment given to the patient and asked to predict the level of improvement in the patient’s health they expect given the treatment provided. A prediction scale was presented below the drug cue ranging from 0 (no improvement) to 100 (full recovery) and participants were asked to make a prediction by clicking on a point on the scale. Clicking on the scale would result in the appearance of a horizontal bar extending from 0 to the final click location. Once a prediction has been made, participants could press the space bar to continue. The patient’s actual observed health improvement was then presented on a near-identical scale below the prediction scale (the only difference being the colour of the horizontal bar inside the scale). Outcomes were animated as a growing horizontal bar along the scale starting at 0 to the final outcome value. Large improvements in health were indicative of the drug’s efficacy in treating the disease. Prediction ratings made across the experiment were our primary measure of learning, and were thus used to determine model fit. In addition to predictions across training, we also measured participants’ summary judgements at the end of the experiment; these include measures of outcome mean, outcome mode and causal ratings judging the efficacy of each treatment on patient recovery.

**Figure 2.**
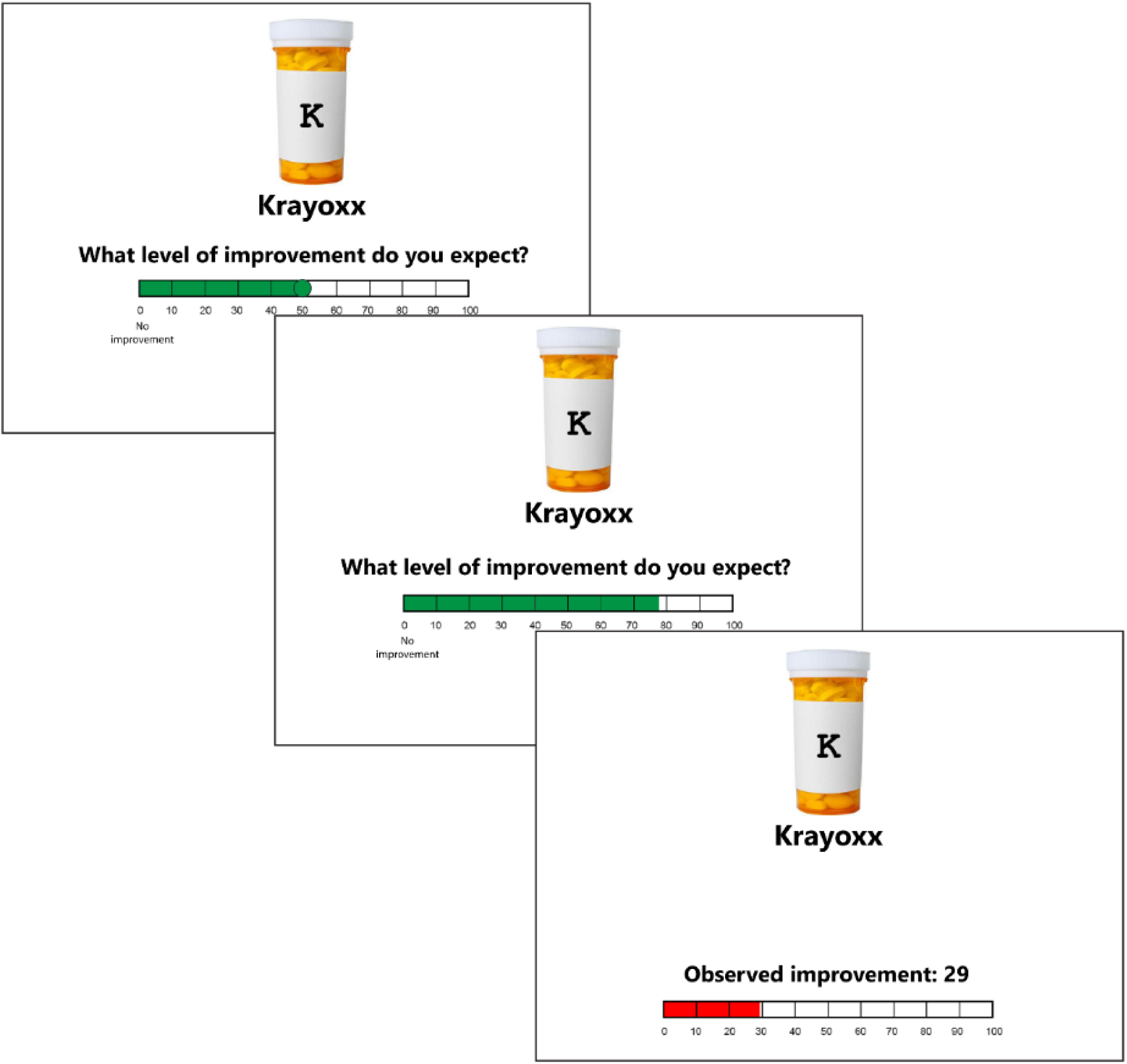
Trial schematic illustrating an example of a single trial sequence in the contingency learning task. Participants were shown a medicine cue and asked to predict the patient’s level of improvement after taking the medicine. Participants could click anywhere on the scale to make their prediction, and press the Continue button to confirm their selection. They were then shown the actual observed health improvement for that patient.

#### Variations across experiments

We present here the rationale for each experiment. More detailed information about the design of each experiment can be found in Supplementary Materials (pp 2-6). Experiment 1 was designed to determine whether participants were sensitive to the presence of variability in outcome values. To achieve this, we presented cues with identical outcome means (mean = 25, 50, 75) but had outcomes that were either constant values (e.g., always 25 on a scale from 0 to 100), or variable values sampled from a uniform or normal distribution. The primary distinction between cues with outcomes from a uniform vs normal distribution lies in the frequency of different outcome values occurring. If participants were sensitive to, and made predictions based on, the experience of variability in outcome values, we might expect differences in their pattern of predictions for cues with constant vs variable outcomes of the same mean. In addition, distribution of predictions is expected to differ for cues with normally distributed vs uniformly distributed outcomes; we expect the Distributed Model to be sensitive to variability in the probability of different outcome values, whereas the Simple Delta approach may fail to discriminate between the two.

In Experiment 2, we manipulated the base-rate of the outcome in a between-subjects design by presenting a number of filler cues, such that the overall outcome base-rate could be low, medium or high (30, 50 or 70) while the mean outcomes of target cues varied across the range of possible outcomes (mean = 15, 30, 50, 75, 80). All target cues had outcomes sampled from a normal distribution. Differences in the base-rate of the outcome have previously been shown to impact judgements people make about predictive cues (e.g., the inverse base-rate effect; Medin & Edelson, 1988; Don et al., 2021) and non-predictive cues (e.g., the outcome density effect in illusory causation; Alloy & Abramson, 1979; Blanco & Matute, 2020; Blanco et al., 2020a). Varying the base rate thus allows us to determine whether contextual factors assert any influence on predictions made to *specific* target cues, and whether these differences, if present, are captured by our two models.

In Experiments 3 & 4, we were interested in whether participants’ ratings were sensitive to the range of probable outcome values. In Experiment 3, we drew outcomes from either a normal, positively-skewed or negatively-skewed distribution at two levels of outcome mean (35 and 65). The logic here is that these distributions differ in the outcome values that are likely to be sampled. For example, one cue A might be associated with a normally distributed outcome with a mean of 65, while another cue B was associated with a positively skewed distribution with the same mean. In the latter case, most of the outcomes shown were above 65 but occasionally outcomes of a lower value were shown. If participants’ judgements reflect the mean outcome to which they are exposed in the presence of that cue, then their judgements for A and B will be equivalent. If participants make judgments that reflect the most likely range of outcomes, then they are likely to predict that the outcome for B will be higher than for A. Alternatively, if participants are more strongly influenced by the rarer but more discrepant low outcomes then they may predict lower outcomes for B than for A.

Experiment 4 provided an additional test of the impact of distribution shape and variance by comparing cues with outcomes sampled from a normal distribution with a small variance (normal-narrow), normal distribution with large variance (normal-wide) and a bimodal distribution. Like in Experiment 3, mean outcome was 35 for half the target cues and 65 for the other half.

### Results & Discussion

The pattern of predictions made for each cue type was compared using Kolmogorov-Smirnov (KS) test of equality on the final 10 presentation of each cue in Experiments 1 & 2, and last 20 presentations in Experiments 3 & 4 (the increase in trials used for analysis in the latter experiments were due to more subtle differences in the distributions being compared). Note that this restriction applied only to empirical analyses—all trials were included in subsequent model fitting. The KS test provides a *D* statistic, which quantifies the difference in the cumulative distribution function of the two comparison distributions; it is therefore a measure of the how similar the two distributions are: larger *D* statistic indicates a larger difference between the two distributions. Using the *Matching* package in R (Sekhon, 2011), a bootstrapping procedure (number of bootstraps = 1000) was applied to calculate a bootstrap p-value for the KS test, with the null hypothesis that probability densities for the two comparison distributions are the same. We assumed a conventional significance threshold of p = .05 for these tests. It should be noted that no corrections were applied for multiple (planned) comparisons however, our aim was not to draw strong inferences from any single finding. Rather, these analyses were intended to provide a descriptive picture of participants’ responses in the task and provide empirical data for model fit. The conclusions of this paper are based on the outcomes of model comparison, rather than on the outcome of any individual statistical test. Overall, the pattern of predictions across all four experiments differed for cues with identical means but with outcome values sampled from different distributions. A summary of the results obtained from comparisons across all four experiments are shown in Table 2.

**Table 2.** Summary of results from Kolmogorov-Smirnov (KS) test of equality. D statistic approaching 1 indicate large differences between distributions. Column values indicate outcome mean and each row is a contrast comparison involving outcome distribution.

| **Experiment 1** |  |  |  |  |  |
| --- | --- | --- | --- | --- | --- |
|  | 25 | 50 | 75 |  |  |
| Fixed vs Variable<br>(average Uniform<br>and Normal) | $D = .160, p = .001$ | $D = .402, p < .001$ | $D = .225, p < .001$ | | |
| Uniform vs Normal | $D = .096, p = .220$ | $D = .154, p = .007$ | $D = .138, p = .021$ | | |
| **Experiment 2** |  |  |  |  |  |
|  | 15 | 30 | 50 | 70 | 85 |
| Base-rate 30 vs<br>Base-rate 50 | $D = .083, p = .560$ | $D = .072, p = .736$ | $D = .133, p = .082$ | $D = .161, p = .019$ | $D = .150, p = .035$ |
| Base-rate 30 vs<br>Base-rate 70 | $D = .126, p = .134$ | $D = .095, p = .427$ | $D = .100, p = .365$ | $D = .156, p = .033$ | $D = .167, p = .017$ |
| Base-rate 50 vs<br>Base-rate 70 | $D = .074, p = .738$ | $D = .128, p = .122$ | $D = .128, p = .122$ | $D = .223, p < .001$ | $D = .218, p = .001$ |
| **Experiment 3** |  |  |  |  |  |
|  | 35 | 65 |  |  |  |
| Normal vs Positive<br>Skew | $D = .085, p = .060$ | $D = .046, p = .694$ | | | |
| Normal vs Negative<br>Skew | $D = .073, p = .156$ | $D = .142, p < .001$ | | | |
| **Experiment 4** |  |  |  |  |  |
|  | 35 | 65 |  |  |  |
| Bimodal vs Normal<br>(average Normal<br>Narrow and Normal<br>Wide) | $D = .083, p = .009$ | $D = .042, p = .491$ | | | |
| Normal Narrow vs<br>Wide | $D = .085, p = .026$ | $D = .073, p = .079$ | | | |

Note. *p*-values reported here are uncorrected for multiple (planned) comparisons.

Analysis of test ratings provided at the end of the experiment showed that ratings were based almost completely on the outcome mean alone, with the exception of small differences when comparing cues associated with the most starkly different distributions (e.g. bimodal vs normal). These results are reported in detail in Supplementary Materials: Empirical Analysis section (pp 6-16). In summary, when asked to provide a single summary rating, participants aggregate their knowledge of the outcome values presented with each cue, and use the mean outcome value to inform their judgements, however predictions for individual trials made across training were sensitive not only to the mean value, but additionally informed by the underlying outcome distribution.

In the following section, we will describe the results from fitting participants’ prediction ratings to the Simple Delta Model and Distributed Model, and provide a formal comparison of the two models in capturing participant behaviour during contingency learning.

### Model fit

For each participant, Bayesian Information Criterion (BIC) was calculated for the Simple Delta Model and the Distributed Model using best fitting parameters; a lower BIC value indicates better model fit. We examined the degree to which the Distributed Model fit the data better than the Simple Delta Model by computing a BIC difference score (ΔBIC) for each participant, ΔBIC = BIC_delta_ – BIC_dist_. A positive ΔBIC indicates better fit by the Distributed Model compared to the Simple Delta Model. Figure 3 is a plot of BIC difference scores for each participant in Experiments 1 - 4. It is noteworthy that almost all participants were better fit by the Distributed Model.

**Figure 3.**
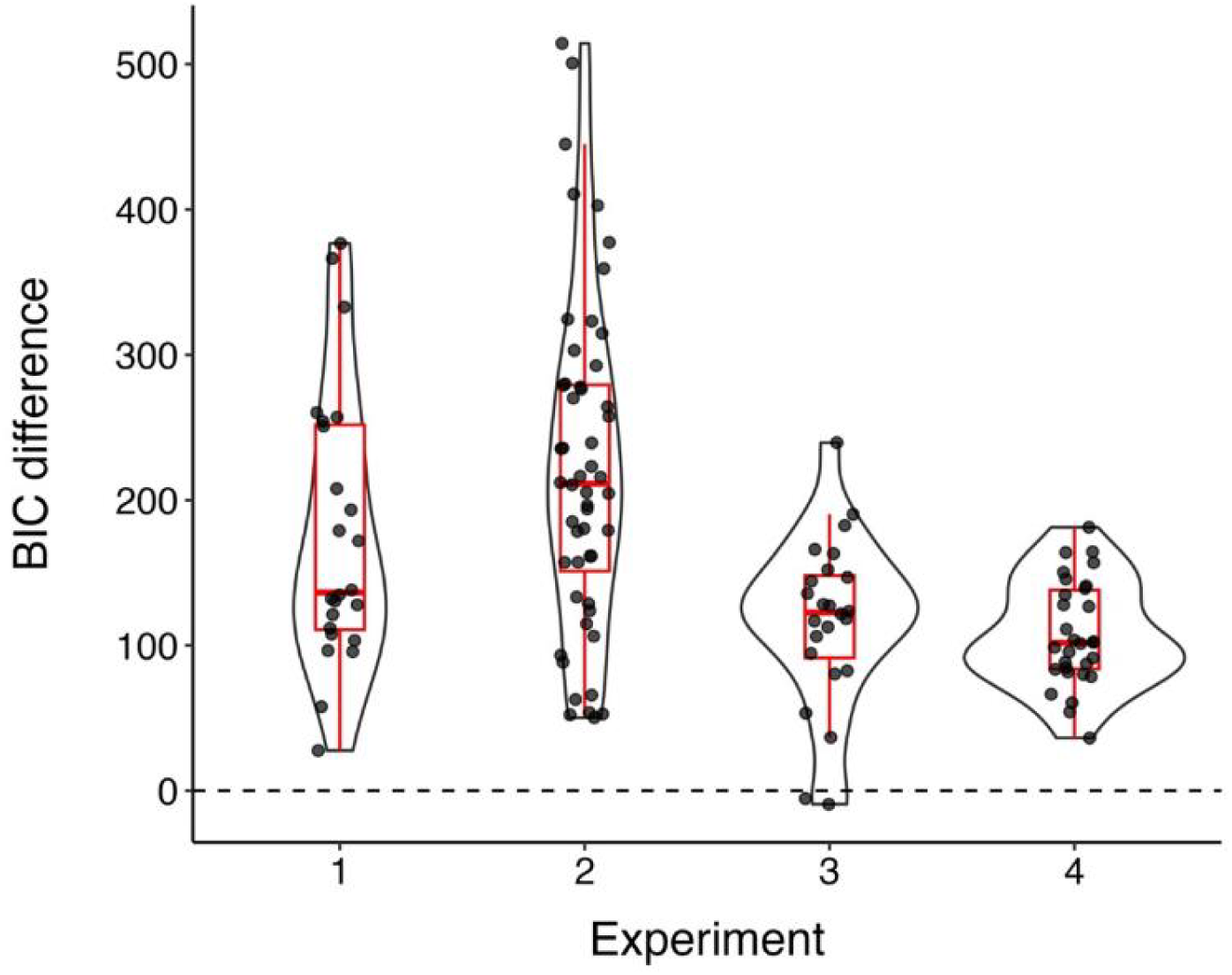
BIC difference score (BIC_delta_ – BIC_dist_) calculated for each participant in Experiments 1 to 4. Positive BIC difference scores indicate better fit by the Distributed Model, whereas negative BIC difference scores suggest better fit by the Simple Delta Model. Boxplots illustrating the median, upper and lower quartile values are shown in red.

Finally, we transformed average ΔBIC for all participants in each experiment into a Bayes Factor, reporting the likelihood ratio of better model fit by the Distributed Model relative to the Simple Delta Model, using the formula described by Wagenmakers (2007): BF_10_ = exp (ΔBIC / 2). Across all four experiments, there is strong evidence in favour of the Distributed Model; BF_10_ > 10 in all experiments (see Table 3). Overall, the Distributed Model provided a better fit of the data compared to the Simple Delta Model. These results suggest that a distributed architecture that enables representation of the underlying distribution of outcome values better captures how human participants are behaving as they make predictions about the outcome during learning. In the following section, we use best fitting parameters for each individual participant to illustrate the terminal associative strengths and discrete predictions generated by both models.

**Table 3.** Bayes Factor in favour of the Distributed model compared to the Simple Delta Model (BF_10_) computed for each experiment. BF_10_ > 10 indicate strong evidence in favour of the Distributed model compared to Simple Delta Model.

| Experiment | $\text{BF}_{10}$ |
| --- | --- |
| 1 | 2.13e+38 |
| 2 | 1.71e+48 |
| 3 | 2.57e+25 |
| 4 | 2.87e+23 |

### Model output

#### Terminal associative strength

The ability for the Distributed Model to represent an associative connection between the cue and each level of the outcome is captured in the terminal associative strengths produced by the model. Unlike the Simple Delta Model which produces a single terminal associative strength, the Distributed Model produces terminal associative strengths linking the cue to each outcome node. This produces a pattern of activation across the dimension that mirrors the underlying distribution (e.g., normal, positive skew, negative skew, bimodal distributions). This is most obvious when associative values are averaged across all participants, as in the inset plots in the top-right corner of each row in Figure 4, but is also demonstrable at the individual participant level (right panel in Figure 4).

**Figure 4.**
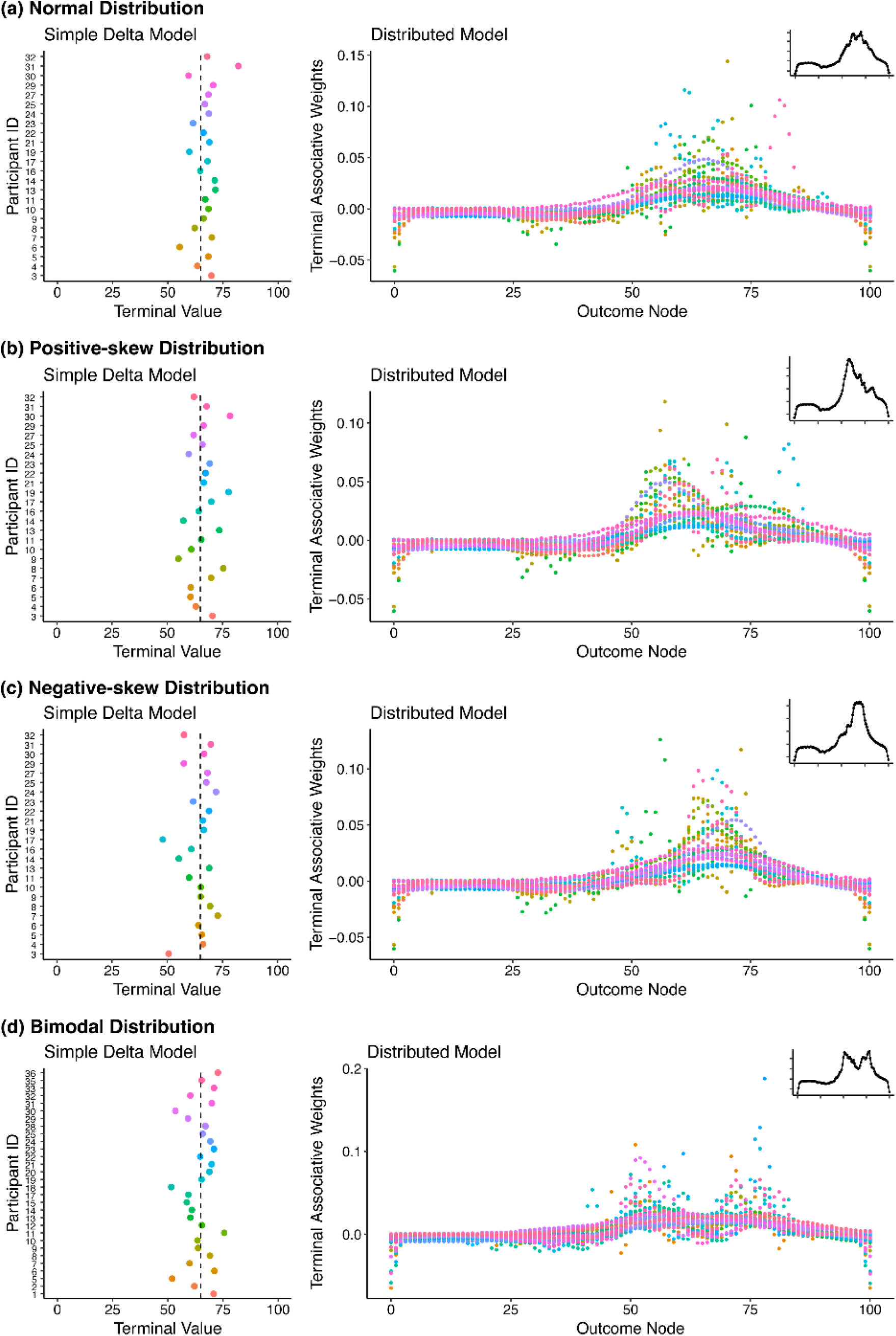
Terminal associative values (ΣV; cue + context) generated by the Simple Delta and Distributed Model for three cues in Experiment 3 and one cue in Experiment 4 with outcome mean of 65, and outcomes sampled from (a) Normal Distribution, (b) Positive-Skew Distribution, (c) Negative-Skew Distribution, and (d) Bimodal Distribution. Terminal values generated by the Simple Delta Model approximates the mean outcome value (vertical dashed line in left panels), whereas the Distributed Model produces associative weights at each outcome node from 0 to 100. *Note*. Different coloured points represent model output for each participant. Inset plot on the top right-hand corner of each Distributed Model output illustrates the average distribution of associative weights across all participants.

Overall, terminal values generated by the model provide some indication that our models are performing as we expected. The Simple Delta Model accurately tracks overall outcome mean but is unable to distinguish between outcomes sampled from different-shaped distributions, whereas the Distributed Model shows sensitivity to the underlying distribution.

#### Simulated model predictions

For both the Simple Delta and Distributed Model, we generated discrete predictions on each trial for every target cue based on Equation 3. This was completed following the trial sequence and outcomes presented to each individual participant, using best-fitting parameters for that participant generated from 100 simulated runs. A critical distinction between the two models when generating discrete predictions is the source of distributional information in the response probabilities. In the Distributed Model, the pattern of associative strengths across outcome nodes naturally provides a distribution of response probabilities used to select a response. In contrast, the Simple Delta Model generates only a single point estimate (approximating the outcome mean) and requires response variability to be introduced artificially by adding Gaussian noise at the point of generating a prediction (as described previously). Nevertheless, we argue that this approach is generous as it provides the Simple Delta Model flexibility to generate variable responses. Thus, while both models use the same response selection process, they differ fundamentally in whether distributional information is learned through the model’s representational architecture (Distributed Model) or assumed through the response generation process (Simple Delta Model).

Figure 5 shows density plots for the 6 target cues presented in Experiment 3, illustrating predictions made by participants (aggregated across all participants) overlayed with simulated predictions generated by the Simple Delta and Distributed model. Visually, there appears to be greater overlap in the distribution of predictions generated by the Distributed model and participants’ actual predictions compared to the Simple Delta model, particularly for cues whose outcomes were sampled from skewed distributions. Predictions generated by the Simple Delta Model appear more diffused around the outcome mean. Figure 6 additionally shows an example of simulated predictions from a *single* participant in Experiment 3, plotted together with the participants’ responses and observed outcome values. The purpose of both figures is to show that the distribution of predictions made by participants were sensitive to both mean and range of observed outcome values, and thus a model that was able to exploit the skewness of the outcome distribution, i.e., the Distributed model, was better able to generate predictions that more closely resembled participant responses.

**Figure 5.**
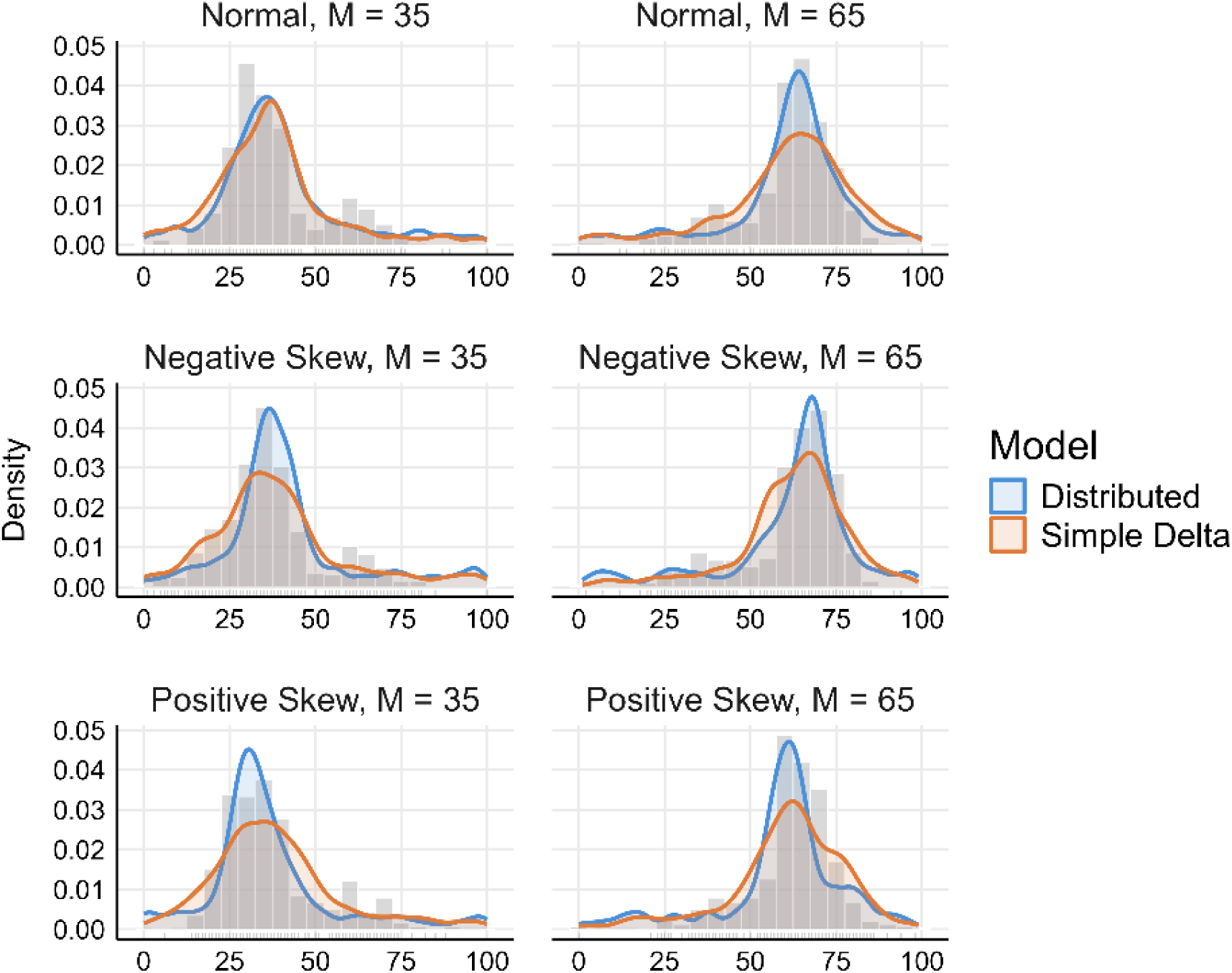
Histogram of participant predictions with overlaid density curve from predictions generated by the Distributed and Simple Delta Model using best-fitting parameters for each participant, for each of the six target cues in Experiment 3.

**Figure 6.**
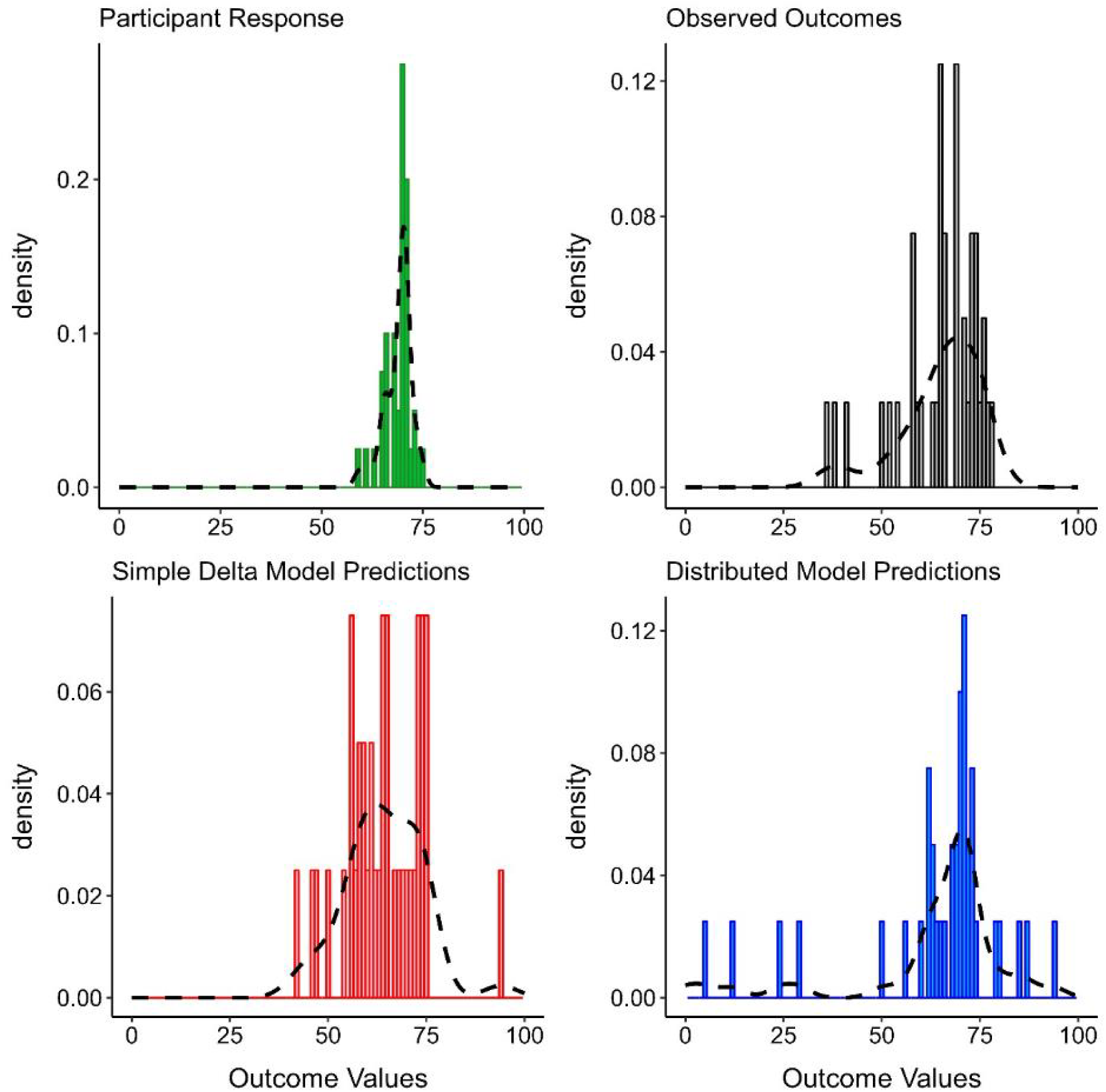
Density plot of (a) participant predictions, (b) observed outcomes, and simulated predictions from the (c) Simple Delta and (d) Distributed Model, for a cue with mean outcome = 65, and outcome values sampled from a negatively-skewed distribution. The illustrated data and model predictions are from a single participant in Experiment 3.

Together, model outputs provide some assurance that the formulation of these models appropriately capture how a simple error-correction model will produce a terminal associative strength that approximates the outcome mean when the outcome has a numeric value, and selective activation of outcome nodes allows for learning about the shape of the underlying outcome distribution. Complete model output from these simulations can be found on the Open Science Framework (Chow et al., 2025). We additionally verified that each model and their parameters were identifiable by simulating data with bounded starting parameter values and assessed the accuracy of the fit to the simulated data (see Supplementary Materials: Computational Modelling Results [pp 17-24]).

## General Discussion

The goal of the present study was to determine whether the behaviour of human participants in a contingency learning task was better captured by a prediction error model capable of representing a distributed network of outcome values, compared to a more conventional class of models that rely on a single point estimate. We first reported empirical data from human participants, and subsequently use participants’ predictions from these experiments to fit the two models of interest. For almost all participants across four experiments, the Distributed Model provided a better fit to the data compared to the Simple Delta Model. Visual inspection of the simulated predictions indicate that the Distributed Model also produced predictions that more closely resembled the pattern of predictions produced by participants in the study. Thus, we show that a model of contingency learning which includes a distributed architecture is better at capturing how humans behave when making trial-by-trial predictions in a contingency learning task when the outcome has continuous values along a dimension.

The results of our experiments are compelling as they suggest that rather than just tracking an overall mean value, participants show some sensitivity to the continuous nature of the outcomes experienced, and are able to use this knowledge (to an extent) when making trial-by-trial predictions about outcomes associated with different cues. One explanation for these findings is that people can engage in distributional learning, and mentally represent outcomes in a probability distribution. This is consistent with a general assumption in cognitive science that people are able to spontaneously acquire knowledge about underlying probability distribution of events and use these learned frequencies to inform behaviour (Griffiths & Tenenbaum, 2006; Chater & Manning, 2006; Lindskog et al., 2021; Schlegelmilch & von Helversen, 2020). However, there is increasing debate about whether people are able to learn and reproduce the structure of underlying distributions: studies that explicitly interrogate distributional knowledge have failed to find evidence of accurate distributional learning when participants were asked to reproduce the distribution of spatial events (Tran et al., 2017), and require extensive guidance (Mason et al., 2025) or constraints to task environment (Brehmer, 1980; Szollosi et al., 2023) in order to be accurate in their reconstruction. A less extreme and perhaps more plausible explanation of the present findings is that participants are not representing complete distributions, but they are sensitive to the frequency of different outcome values occurring for each cue type. This allows them to develop strategies or rules to distinguish between the cues, reflected in the differences in their pattern of predictions. This latter account is no less impressive, given the complexity of the present experiments where participants were presented with up to 13 different cues (intermixed) that differ in overall mean and outcome distribution. Under this account, participants are, at the very least, building sufficiently distinct representations about each cue-outcome relationship to allow them to make predictions that differ from a similar cue whose outcomes are sampled from another distribution (see also Mason et al., 2025). Notably, the Distributed Model captures this sensitivity without explicitly representing distributional parameters such as variance or shape—distributional sensitivity emerges instead from the accumulation of associative weights across outcome nodes, applying the same basic delta-rule learning mechanism across a distributed outcome representation—and without retaining individual outcome episodes in memory as a sample-based account would require. That this architecture is sufficient to account for human sensitivity to different distributions raises the question of whether and when more explicit representational architectures, such as those that track distributional parameters directly or approximate distributions from stored exemplars, are necessary to explain human distributional learning.

One alternate explanation of our findings is that participants are not learning about the outcomes associated with each cue *at all*, but are instead engaging in strategic matching behaviour by making predictions that match the outcome value experienced on the previous trial for that cue type. To account for this possibility, we compared the likelihood of such a “matching model” –using the outcome on trial *n* as a predictor of participants’ predictions on trial *n+1* for a given cue—to a “non-matching model”, where responses are based on the average observed outcome up to trial *n* (i.e., learning). Overall, we found no convincing evidence that participants were systematically engaging in such a matching strategy (see Supplementary Materials: Analysis of Matching Strategy pp 24-26). Indeed, when we correlated individual participant predictions made on trial *n+1* to the observed outcome on trial *n*, we failed to find systematic evidence of a statistically significant correlation across participants and cue type, suggesting that participants’ pattern of predictions cannot be fully explained by a simple matching strategy. A visual summary of this analysis is presented in Figure 7; we illustrate only the correlations from Experiment 3 where there may be increased incentive to use a matching strategy due to greater similarity in the outcome distributions. We find that most participants do not systematically engage in matching behaviour when making predictions. A similar analysis comparing model predictions generated based on best-fitting parameters for the Simple Delta and Distributed Model for each individual participant showed that the Simple Delta Model substantially over-estimates the positive correlation, generating predictions that resemble matching behaviour significantly more so than participants’ actual responses (see Figure H in Supplementary Materials p 26); this pattern was not observed for predictions generated by the Distributed Model.

**Figure 7.**
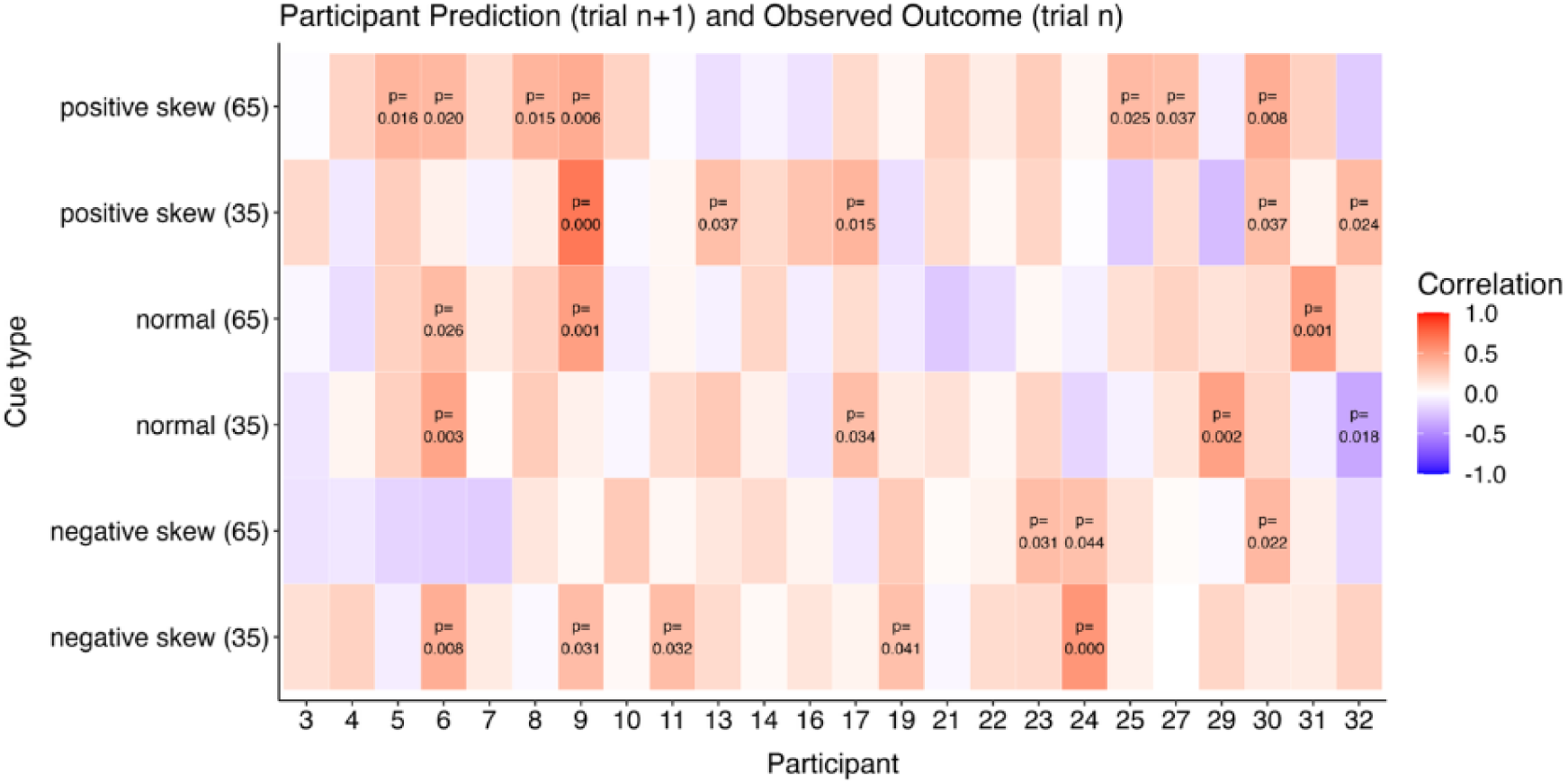
Correlation matrix illustrating the strength of the correlation between participants’ predictions on trial n+1 and the observed outcome on trial n for each cue type in Experiment 3. Each column represents data from an individual participant denoted by their unique participant ID on the x-axis, and each row consists of one of the target cues presented in the study; row labels indicate the distribution type with the outcome mean in parentheses. Cells that appear more red in colour indicate increasingly positive correlation values, whereas bluer cells indicate increasingly negative correlation values, with colours close to white indicating little to no correlation. P-values for statistically significant correlations are included in the relevant cells, whereas correlations which did not reach statistical significance are not shown for visual clarity.

Taken together, we propose that when participants were presented with continuous outcomes in a contingency learning task, their pattern of predictions across learning goes beyond matching the outcome observed on the previous trial and a simple learning strategy that tracks the overall mean. Results from model fit and model comparison additionally provide good evidence in support of the idea that participants’ behaviours are not simply a result of ‘noisy’ responding, but reflect a more sophisticated learning process that allows them to distinguish between similar cues by leveraging features of the underlying outcome distribution.

### Measurements of learning

To our knowledge, the present experiments are the first to investigate learning about continuous events in a contingency learning experiment using predictions made *during* learning. In contrast, studies focused on eliciting distributional knowledge have focused on participants’ ability to accurately reproduce the distribution of experienced events after the fact (e.g., Tran et al., 2017; Mason et al., 2025). Causal learning studies similarly rely on judgements made at the end of the experiment to determine what participants have learned about the cue-outcome relationship (e.g., Shanks & Dickinson, 1988; Wasserman et al., 1993; Vadillo et al., 2005). The appeal of measuring participants’ learning at the end of the experiment is clear: it provides an opportunity for participants to integrate and report the acquired knowledge without interference from ongoing learning processes. This is pertinent in circumstances where the goal of the study is to determine *what* participants have learned (e.g., underlying task structure in distributional learning or the cue-outcome contingency in causal learning experiments), rather than *how* participants may be learning. Indeed, post-experimental estimation tests are often limited as they do not require participants to reproduce the distribution of events, but rather encourage integration across the entire experiment (Collins & Shanks, 2002). In the present experiments, we similarly find an insensitivity of test ratings (average outcome, outcome mode, causal ratings) to the shape of the underlying distribution, despite differences in the pattern of predictions made during training.

In the present study, outcome representation is inferred through the predictions made by participants for different cues across the learning phase: we present participants between 8 and 13 different cues in an intermixed fashion, and they are required to make an outcome prediction on each trial. Participants were never told that they were being assessed on their distributional knowledge, and they were not constrained in how they could respond on any given trial—participants could, in theory, report the outcome mean on every trial, resembling predictions by the Simple Delta Model—thus eliminating any biases that might emerge from using a more explicit test of distributional knowledge, such as strong priors about normality and unimodality, as well as allowing for the influence of specific semantic knowledge like the proportion of people who benefit or do not benefit from treatment. In addition, the task is complex as it involves tracking multiple cues simultaneously. This complexity requires participants to use the outcomes to discriminate between different cues. This feature of the design may be pertinent for providing the necessary scaffolding for participants to learn the structure of the task environment: participants’ attention may be drawn towards differences in outcome values associated with each cue in order to distinguish between them, thus encouraging representations that are sensitive to that distinction. This possibility is consistent with the idea that highlighting aspects of the task important for successful distributional learning increases performance accuracy (Szollosi & Newell, 2020), and evidence from the multi-tasking literature showing greater separation of task representations when two tasks were performed in an intermixed fashion (Garner & Dux, 2015).

Although the use of prediction responses is well suited for our present purposes, we are cautious not to overclaim the benefit of this method for studying distributional learning *per se*, since participants’ pattern of predictions were still significantly different from the actual observed outcome distribution (see Tables A-D in Supplementary Materials: Empirical Analysis), suggesting imperfect distributional knowledge despite the task design encouraging discrimination between cues. Nevertheless, the present method of measuring outcome representation is well suited to test the Distributed Model against a Simple Delta Model, and provides a good test bed for determining whether people are capable of more sophisticated means of encoding outcomes in a contingency learning task with continuous outcomes. To this end, the present findings suggest there is much to be gained from focusing on the pattern of predictions made across the course of learning. These predictions capture differences that are lost when asked to provide a single point estimate at test, at least when the research question is focused on understanding how participants are learning in a noisy and complex environment.

### Applications to human contingency learning

We have shown that participants’ predictions across learning are not merely a result of noisy approximations of observed outcomes, but instead reflect a degree of sophistication in encoding that preserves their continuous and distributed nature. Consequently, a model capable of representing continuous outcomes in a distributed fashion provides a more accurate representation of how participants engage with and learn about continuous outcomes during contingency learning. A notable feature of the Distributed Model approach is its ability to capture variability in responding via distributed representation of the properties of each outcome, rather than via symbolic estimation of uncertainty or explicit computation of sample variance. We do not claim that this approach is computationally simpler than parametric alternatives; indeed, maintaining associative strengths across a large set of outcome nodes is itself non-trivial. Rather, our argument is a mechanistic one: sensitivity to distributional shape need not arise from a dedicated cognitive mechanism that explicitly tracks uncertainty, but can instead emerge from the pattern of associations distributed across outcome nodes, with the frequency of experienced outcomes encoded in the strength of these associations. Many models, particularly Bayesian approaches, require researchers to specify parametric representations of uncertainty (e.g., variational approximations of posterior distributions), which may introduce arbitrary constraints or assumptions about how individuals perceive and respond to variability. In contrast, the Distributed Model allows for the emergence of variable responding through associative structure alone.

This mechanistic approach has several advantages. First, it connects directly to the extensive associative learning literature that has successfully modelled phenomena across species using similar connectionist frameworks (e.g., Blough, 1975; McLaren & Mackintosh, 2000, 2002; Ghirlanda & Enquist, 1998; Livesey & McLaren, 2019). Second, it suggests that the capacity to respond to uncertainty emerges naturally when there is sufficient complexity in how the model represents stimuli. While there are several approaches to modelling variation in predictions and the effect of uncertainty on learning, this is typically achieved by assuming a distinct cognitive mechanism that computes and updates explicit uncertainty. Our approach suggests that even if symbolic representation of uncertainty is sufficient, it is not necessary to capture learning about variable outcomes in the conditions tested here. Models with formal calculations of uncertainty and variance provide a powerful and often convenient way to model learning processes, and may well help to characterise certain aspects of human cognition accurately. However, more research is needed to investigate precisely what types of learning phenomena necessitate this approach.

So far, we have presented two strengths of the Distributed Model for modelling contingency learning with continuous outcomes: first, it is a better model of what people are actually learning in these tasks compared to a Simple Delta Model, and second, it does not require additional assumptions about the role of uncertainty in learning. A third notable advantage of the distributional approach is that it presents a useful tool for understanding the learning processes underlying (biased) causal inference. This third advantage becomes particularly evident when considering how people reason from continuous rather than discrete events. The potential for ambiguity when interpreting continuous outcomes creates more opportunities for cognitive biases to shape both what is learned and the causal beliefs that follow.

The ability for people to learn about continuous outcomes is largely ignored in the contingency learning literature. Most prominent theories of causal learning are based on the assumption that people represent their experiences in a discrete way, where the cue and outcomes are either present or absent. The frequency of these discrete events (e.g., patient takes a medicine [cue present] and recovers from their illness [outcome present]) are subsequently used to form a causal judgement (e.g., delta p metric; Allan, 1980). Indeed, one interpretation of our findings is to assume that the different pattern of predictions generated for different cues are simply a product of participants applying *discrete* categorisation rules without appealing to the shape of the distribution—e.g., treatment A tends to have outcomes around 50 but sometimes higher than 50. Nevertheless, existing covariation-based models largely ignore the question of *how* noisy, continuous events in the real world are categorised into discrete categories (Ahn et al., 2007; Marsh & Ahn, 2009). Take for example, the scenario used in the present studies. Participants were told that high-value outcomes indicate greater patient recovery, with the extreme ends of the scale indicating no recovery (outcome = 0) or full recovery (outcome = 100). When participants experience extremely high or low outcome values, categorisation of the outcome into patient recovered or not recovered may seem obvious, however what is less obvious is how to classify an outcome value around the middle of the scale (e.g., 50/100). These outcomes are ambiguous relative to the goals of the learner, and may present a challenge to contingency models that rely on discrete categorisation of events for causal inference.

We propose that the interpretation and classification of continuous and potentially ambiguous events in learning is a flexible process shaped by several factors, including the underlying probability distribution and the learner’s prior beliefs. Future development of the Distributed Model could incorporate mechanisms for representing prior beliefs and exploring how these beliefs interact with distributional learning. For example, if a learner has strong prior beliefs about the efficacy of a treatment producing recovery, they might encode outcome values differently depending on whether the treatment was administered, thus biasing their representation of the outcome distribution in ways that reinforce existing beliefs. Variability in learning through this mechanism may be especially pronounced under zero contingency scenarios, where there is no objective causal relationship between cue and outcome. It has previously been shown that when participants are uncertain about the causal status of a cue, like in zero contingency scenarios, there is a tendency to “explain away” conflicting information in ways that allow them to preserve their pre-existing beliefs (Chow et al., 2024; Livesey et al., 2025; Spicer et al., 2020; Spicer et al., 2022). Testing such a model – incorporating both prior beliefs and a distributed representation – would require explicit computational mechanisms that are not currently present in the Distributed Model. Nevertheless, such extensions are possible and could provide insights into the formation and maintenance of false beliefs in real-world scenarios with variable and noisy outcomes. This possibility presents an important direction for future research.

### Conclusions

In summary, we have shown that when presented with a continuous outcome, participants in a contingency learning task produced a pattern of responding that suggested sensitivity to the underlying outcome distribution. We modelled the results from four empirical experiments and found that model fits were significantly better for virtually all participants using a Distributed Model which has the capacity to represent a network of associative weights across the entire outcome dimension, compared to a Simple Delta Model which only tracks the overall mean. Contrary to most conventional models of learning that simplifies the content of what is learned, these findings suggest that variability in responding is not merely a product of noise, but that people are capable of more sophisticated encoding processes that preserve their continuous and variable nature. These findings help advance our understanding of how people learn from experienced events in a complex and noisy environment, and provides a framework for thinking about causal inference in the real world.

## Supporting Information

**S1 Supplementary Materials**. Includes analysis of empirical data for each experiment, and model recovery and parameter recovery analysis.

