## Supplementary material for "Predicting continuous outcomes: Some new tests of associative approaches to contingency learning": S1 Supplementary Materials

**to accompany**

Julie Y. L. Chow^1^

Hilary J. Don^2,3^

Ben Colagiuri^3^

Evan J. Livesey^3^

*^1^School of Psychology, UNSW, Sydney*

*^2^ School of Psychology, University College London*

*^3^ School of Psychology, The University of Sydney*

**Empirical Analysis**

Here we report the empirical data from four separate contingency learning experiments. The primary goal of these experiments was to obtain participant data for model fitting: we were interested in whether response patterns produced by participants when predicting what outcomes will follow from a specific cue was better captured by the Distributed Model that allows for distributional representation of continuous outcome values – that is, whether there is any evidence that human participants show sensitivity to the underlying distribution the outcomes are sampled from –or whether their behaviour reflects tracking of the overall mean and some variance around the mean estimate, captured in the Simple Delta Model. Note that our focus is not on revealing whether participants *do* accurately represent these distributions during learning, but instead on whether there is any indication in their pattern of predictions of a *sensitivity* towards differences in the underlying outcome distribution (and the frequency or types of values they produce) when making predictions for cues that predict a fixed vs variable outcome (Experiment 1), when the underlying outcome base-rate is different (Experiment 2), and when the outcomes are sampled from different-shape distributions (Experiments 3 & 4).

**Method**

**Participants**

All participants were undergraduate students at the University of Sydney enrolled in an introductory psychology course. Participants were awarded partial course credit for their participation. Initial sample size for each study were as follows: Experiment 1, *n* = 28; Experiment 2, *n* = 52 (Base-rate 30: *n* =18, Base-rate 50: *n* =18, Base-rate 70: *n* =16); Experiment 3, *n* = 32; Experiment 4, *n* = 36.

**Stimuli & Apparatus**

The experiment was programmed using MATLAB and the Psychophysics Toolbox extensions (Brainard, 1997; Pelli, 1997). Cue stimuli were presented at the top of the screen in the form of a pill bottle with different letters A-M on them presented together with the name of the fictitious drug underneath the pill bottle. Drug names for cues A-M are as follows (in order): Ambrosia, Blaccine, Cloveritol, Dioxnyl, Ephemerol, Felicium, Gambutrol, Hyronalin, Imobatine, Jamitol, Krayoxx, Lithorol, Metazine.

**Design**

In Experiment 1, participants experienced outcomes with a mean value of 25, 50 or 75. The outcome values presented to participants on any given trial for the different cue types were either a fixed value (e.g., always 25 on a scale from 0-100) or sampled from a normal or uniform distribution. For outcomes sampled from the variable distributions, the set of outcome values experienced were identical across the two distributions at each level of outcome mean, but differed in the frequency of those values occurring throughout the study (see Figure A). For example, for a distribution with a mean of 50, the different outcomes presented were values between 30 and 70 in 5-point increments for both the normal and uniform distribution. However, the relative frequency in which these values were presented differed, with 50 being the most common outcome for the normal distribution, whereas all values were equally likely in the uniform distribution.

**Figure A.**

*Number of trials with specific outcome values presented to participants in Experiment 1 as a function of distribution type (fixed value, normal, uniform distribution) and outcome mean (25, 50, 75).*

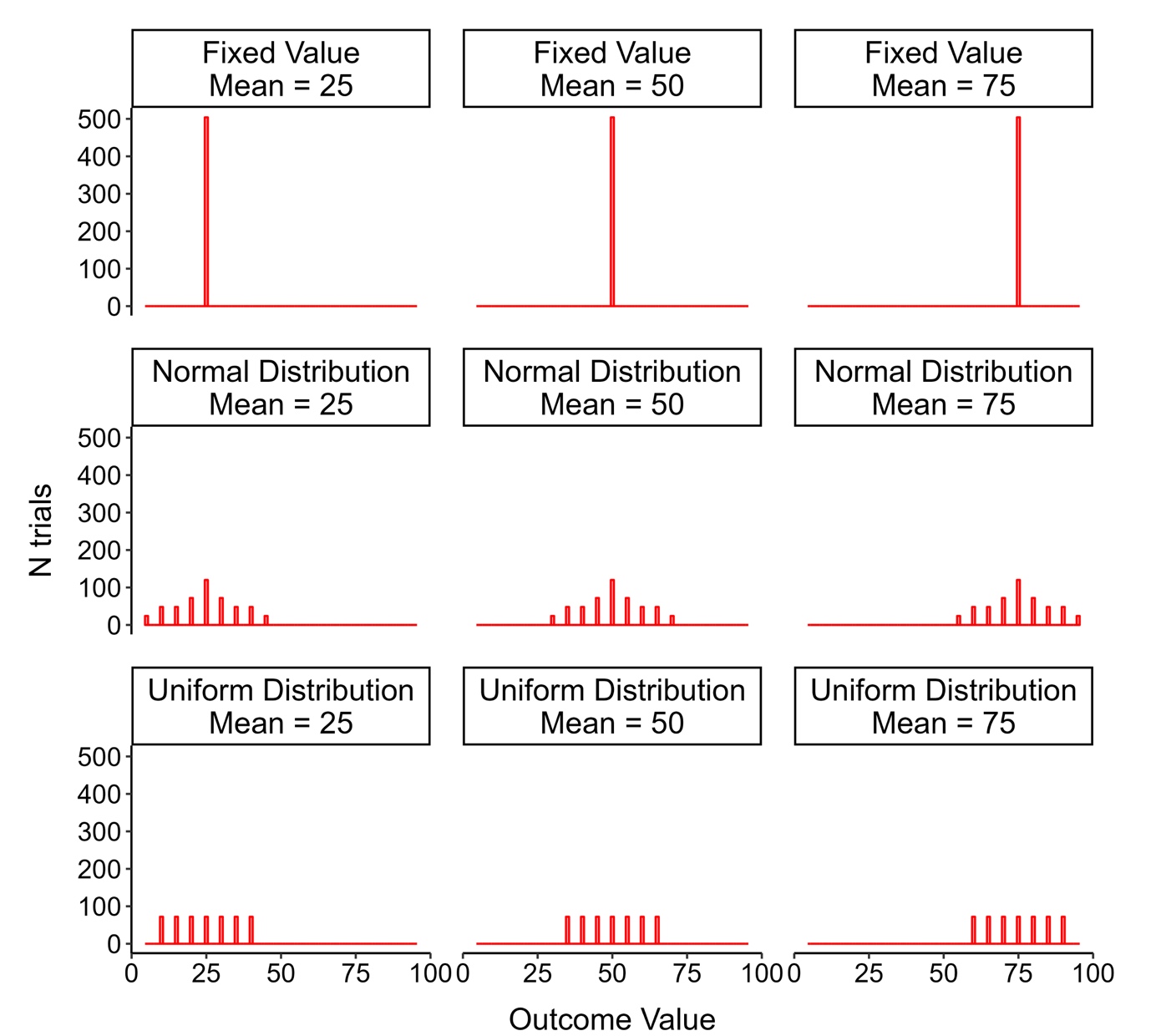

Experiment 2 used a between-subjects design to manipulate the base-rate of the outcome, such that the overall base-rate was 30, 50 or 70; this was achieved by presenting six filler cues with outcomes close to the intended base-rate value. In the base-rate 30 condition, filler cues had outcomes sampled from normal distributions with means = 15, 17, 18, 19, 20 and 21. In the base-rate 50 condition, filler cue means = 47, 48, 49, 51, 52, 53, and finally in the base-rate 70 condition, filler cue means = 79, 80, 81, 82, 83, 85 (all σ = 5, range = ±15). All target cues had outcomes sampled from a normal distribution, with a mean value of 15, 30, 50, 70 or 85. In this and subsequent experiments, we also included a test phase where participants were asked to report the *average* outcome for each of the 5 target cues.

Experiments 3 & 4 directly manipulated properties of the distribution. In Experiment 3, participants saw outcomes sampled from either a normal, positively-skewed or negatively-skewed distribution with an outcome mean of 35 or 65. Outcome values were generated separately for each cue from distributions at each level of outcome mean. For normally distributed outcomes, samples were drawn from a truncated Gaussian distribution with σ = 9.2 and values restricted to lie within ±34 points from the mean. This ensures that outcome values stayed within a specified range around the mean. For skewed outcomes, values were sampled from an ex-Gaussian distribution with σ = 3, where the Gaussian location parameter μ was adjusted (e.g., μ = 25 for positively skewed distribution with outcome mean = 35) to ensure that after applying skew using the parameter τ (τ = +10 for positive skew, τ = −10 for negative skew), the final sampled distribution had the desired mean (35 or 65). The ex-Gaussian parameters were selected so that the resulting distribution had an overall mean of either 35 or 65, while maintaining the same ±34 range as the normally distributed outcomes.

In Experiment 4, we compared normally distributed outcomes with high (normal-wide) or low variance (normal-narrow) against a bimodal outcome distribution. Bimodal distributions comprise of a concatenation of two normal distributions: a bimodal distribution with outcome mean of 65 comprised of the concatenation of a normal distribution centered on a mean of 55 (range = ±24 points) and a normal distribution centered on a mean of 75 (range = ±24 points), whereas a bimodal distribution with outcome mean of 35 comprised of a normal distribution with mean of 25 and 45 (range = ±24 points from the mean). Normal distributions with the same outcome mean were created to match the standard deviation present in the bimodal distribution either by matching the variance of one of the two components of the bimodal distribution (normal narrow distribution, σ = 5, range = ±34 points), or by matching the standard deviation of the bimodal distribution as a whole (normal wide distribution, σ = 11.35, range = ±34 points). Like in Experiment 3, mean outcome was 35 for half the target cues and 65 for the other half. In addition to average outcome estimation at test, we also included two measures of the most likely outcome (outcome mode) and a causal rating of the efficacy of each target drug in treating the disease. Efficacy ratings were made on a scale from -100 (Effectively worsens the disease) to 100 (Effectively treats the disease).

**Results**

Our primary analysis was to compare the distribution of outcome predictions made for cues sharing the same outcome mean but with outcomes sampled from different distributions. These results are reported in the main text of this paper (Table 2). Here, we present additional analyses comparing the pattern of predictions produced by participants in the final half of their observations to the *observed outcomes* for each cue type. The idea here is that if participants were accurately representing the entire distribution outcomes were sampled from, their pattern of predictions should in theory not systematically differ from that underlying distribution. However, as noted in the Introduction of the main text, we do not anticipate such complete representation of distributional knowledge, and particularly not in situations where participants only experienced up to 40 presentations for each cue type. In Experiments 2 – 4, we additionally included test measures at the end of the experiment that required participants to estimate the *average* outcome associated with each cue (Experiment 2 – 4), the *most frequent* outcome (Experiments 3 & 4) and to provide a causal rating for the efficacy of the treatment in curing patient illness (Experiments 3 & 4).

In all experiments, participants saw two treatments that always resulted in a fixed outcome of 0 (fixed-0) and 100 (fixed-100). These cues served as attention check cues to ensure only participants who were engaged in the task throughout the learning phase were included. Participant data was excluded (from statistical analyses and model fitting) if mean prediction ratings for fixed-0 and fixed-100 were greater than 10 or less than 90 respectively, averaging across the final 10 (Experiments 1 & 2) or final 20 trials (Experiments 3 & 4). We increased the number of trials used to compute the exclusions in latter experiments due to the more subtle differences in outcome distributions. Final dataset after exclusions were: Experiment 1 = 24; Experiment 2 = 52 (across two conditions); Experiment 3 = 24; Experiment 4 = 30.

**Experiment 1**

***Comparing predictions to observed outcomes***

Comparison of predictions to observed outcomes for each cue in the second half of presentations (10 presentations of each cue type) revealed statistically significant differences for almost all cue types in the pattern of predictions compared to the underlying outcome distribution (see Table A). This is not particularly surprising given the difficulty of the task with intermixing cue presentations, and the limited number of experiences with each cue type.

**Table A**

*D statistic and p-value from the Kolmogorov-Smirnov test of equality comparing the distribution of predictions to the observed outcomes across the last 10 trials, as a function of distribution condition (fixed value, uniform, normal distribution) and outcome mean (25, 50, 75), average across all participants.*

|  | 25 | 50 | 75 |
| --- | --- | --- | --- |
| Fixed | *D* = .413, *p* < .001 | *D* = .096, *p* = .220 | *D* = .350, *p* < .001 |
| Uniform | *D* = .225, *p* < .001 | *D* = .254, *p* < .001 | *D* = .208, *p* < .001 |
| Normal | *D* = .171, *p* = .002 | *D* = .213, *p* < .001 | *D* = .225, *p* < .001 |

**Experiment 2**

In Experiment 2, we tested whether predictions for target cues with outcomes sampled from identical normal distributions were affected by differences in the overall base-rate. This was a between-subjects design where participants experienced an outcome base-rate of 30 (n = 18), 50 (n = 18) or 70 (n = 16) generated by the presence of filler cues. Importantly, target cues were sampled from identical normal distributions with a mean of 15, 30, 50, 70 and 85 respectively.

***Comparing predictions to observed outcomes***

Table B summarises the results from KS test comparing pattern of predictions to the observed outcomes in the final half of the study (13 presentations per cue type).

**Table B**

| Mean | Base-Rate 30 | Base-Rate 50 | Base-Rate 70 |
| --- | --- | --- | --- |
| 15 | *D* = .200, *p* < .001 | *D* = .170, *p* = .001 | *D* = .159, *p* = .005 |
| 30 | *D* = .133, *p* = .016 | *D* = .196, *p* < .001 | *D* = .121, *p* = .060 |
| 50 | *D* = .144, *p* = .007 | *D* = .119, *p* = .045 | *D* = .196, *p* < .001 |
| 70 | *D* = .093, *p* = .197 | *D* = .226, *p* < .001 | *D* = .229, *p* < .001 |
| 85 | *D* = .163, *p* = .002 | *D* = .126, *p* = .028 | *D* = .329, *p* < .001 |

***Test Ratings***

In this experiment, we also asked participants to report the *average* outcome expected for each of the target cues. Mixed-model ANOVA comparing average prediction for each cue as a function of base-rate group showed a main effect of cue type on mean estimates, *F*(3.1, 152.1) = 984.9, *p* < .001, η_p_^2^ = .953, no significant effect of base-rate group, *F*(2,49) = .179, *p* = .836, η_p_^2^ = .007, and no interaction between cue type and group, *F*(6.21, 152.1) = 1.58, *p* = .155, η_p_^2^ = .060. Overall, all participants were highly accurate at predicting the mean outcome value, with *t*-tests revealing no significant difference in participants’ average predictions and the actual mean, all *p* > .05. These results are depicted in Figure B.

**Figure B**

*Average outcome estimates (large white circle ± SEM) made at test in Experiment 2 for each of the five target cues (mean = 15,30,50,70,85) grouped by base-rate condition (30, 50, 70). Each open circle with coloured outline represents an individual participant data point plotted over boxplots illustrating the inter-quartile range and median for each condition.*

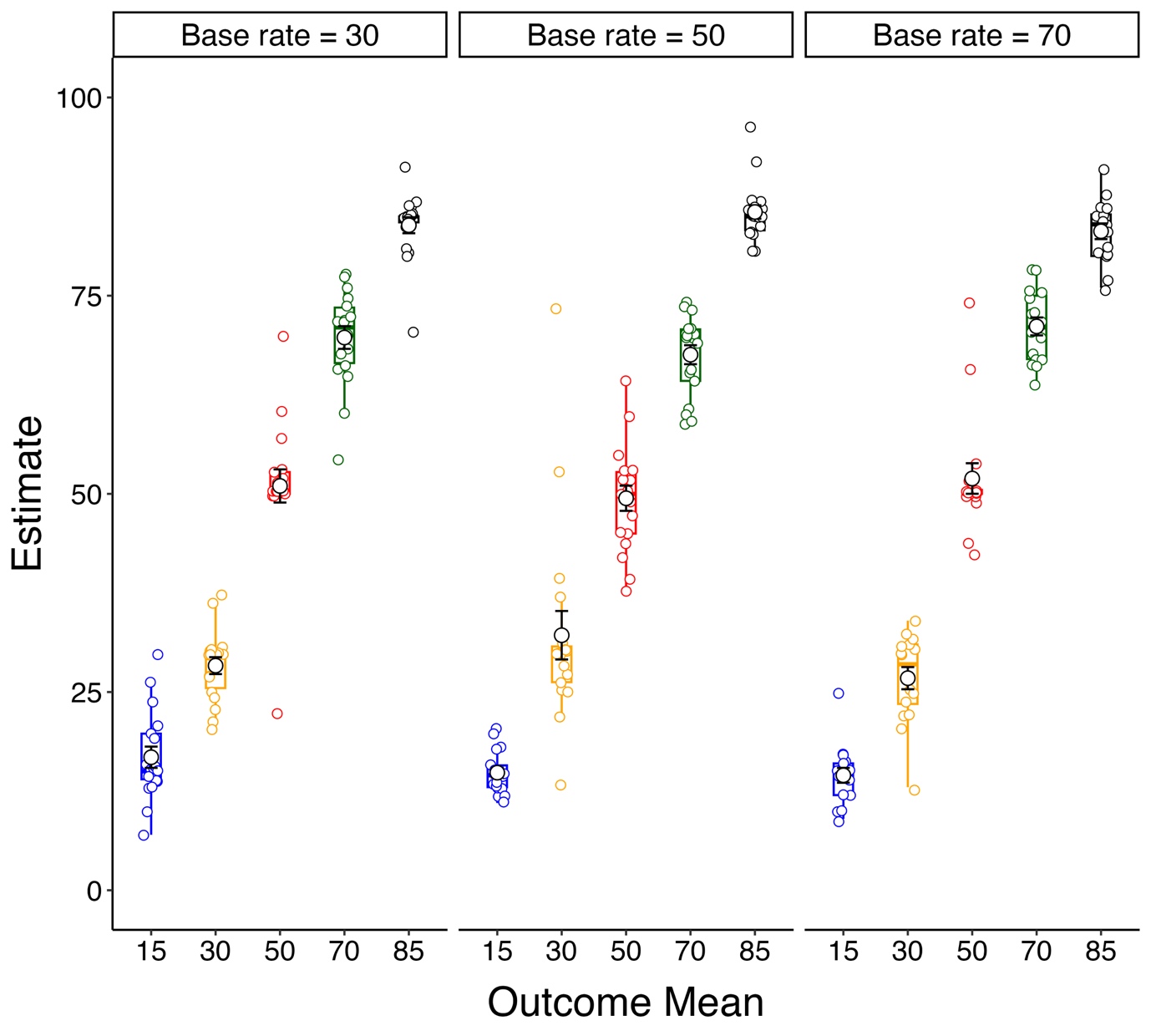

**Experiment 3**

Experiment 3 was designed to determine whether the pattern of predictions made by participants were different for outcomes sampled from a normal distribution compared to a positively skewed and negatively skewed distribution with the same outcome mean. Note that unlike Experiments 1 & 2, we analysed ratings from the last 20 presentations of each cue type in Experiments 3 & 4 due to the more subtle differences between distributions.

***Comparing predictions to observed outcomes***

A comparison of participants’ predictions to the observed outcomes revealed significant differences in the distribution of values across all levels of outcome mean and distribution type (Table C).

**Table C**

*D statistic and bootstrap p value obtained from a Kolmogorov-Smirnov test of equality comparing prediction ratings to observed outcomes in the last 20 presentations of each cue type in Experiment 3.*

|  | 35 | 65 |
| --- | --- | --- |
| Normal | *D* = .146, *p* < .001 | *D* = .158, *p <* .001 |
| Negative Skew | *D* = .158, *p* < .001 | *D* = .115, *p* *=* .004 |
| Positive Skew | *D* = .121, *p* = .002 | *D* = .146, *p* *<* .001 |

***Test Ratings***

Like in Experiment 2, we asked participants to predict the average outcome for each of the six target cues. Results from the test phase are shown in Figure C. Given our interest in comparing only the independent effects of outcome mean and distribution shape on participants’ mean estimates (and not the interaction between the two), we ran three planned contrasts comparing 1) normal and positive-skew distribution, 2) normal and negative-skew distribution, 3) mean 35 and mean 65. Here, we found only a significant main effect of outcome mean, *F*(1,23) = 311.7, *p* < .001, η_p_^2^ = .931, with significantly higher mean estimates for cues with outcome mean of 65 compared to outcome mean of 35. There was no statistical difference in mean estimates when comparing cues with outcomes sampled from a normal vs negatively-skewed distribution, *F*(1,23) = .854, *p* = .365, η_p_^2^ = .036, and when comparing outcomes sampled from a normal vs positively-skewed distribution, *F*(1,23) = .134, *p* = .717, η_p_^2^ = .006. For all six cues, participants’ mean estimates were close to the objective mean outcome value, suggesting high accuracy at tracking the outcome mean.

In addition to mean estimates, we also asked participants to report the most frequent outcome for each of the target cues. This measure was included to help us determine whether participants were sensitive to range of probable values within the distribution, whereby outcomes sampled from a positively skewed distribution will have more values greater than the mean, and fewer values below the mean, and this is reversed for a negatively skewed distribution with the same mean value. If participants’ modal estimates were influenced by the shape of the distribution, we might expect estimates to be lowest for the negative-skew distribution (most outcomes lower than the mean) and highest for the positive-skew distribution (most outcomes higher than the mean), with normal distribution in between, for each level of outcome mean. Planned contrast comparisons again found participants’ ratings to approximate the mean: there was a significant main effect of outcome mean, *F*(1,23) = 237.8, *p* < .001, η_p_^2^ = .912, but no significant difference in estimates for our two distribution contrasts, *Fs*(1,23)$\leq$ 1.18, *ps* $\geq$ .288, η_p_^2^ $\leq$.05.

Finally, participants’ judgements about the efficacy of each of the target treatment cues revealed only a main effect of outcome mean, *F*(1, 23) = 42.2, *p* < .001, η_p_^2^ = .647, with greater ratings for cues with outcome mean = 65 (M = 42.5, SD = 27.1) than cues with outcome mean = 35 (*M* = 3.32, SD = 32.0). There were no differences in efficacy ratings as a function of outcome distribution, *Fs*(1,23) $\leq$ 2.83, *p*s $\geq$ .106, η_p_^2^ $\leq$ .110. Overall causal ratings made at the end of training appear to be sensitive to the mean outcome but agnostic to manipulations of outcome distribution.

**Figure C**

*Average ratings at test in Experiment 3 (mean denoted by white circle ± standard error), when asked to select (a) the average outcome value, (b) the most common outcome value associated, and (c) efficacy of the drug at treating the illness, separated by outcome mean (35, 65) and outcome distribution (Normal, Positive-skew, Negative-skew).*

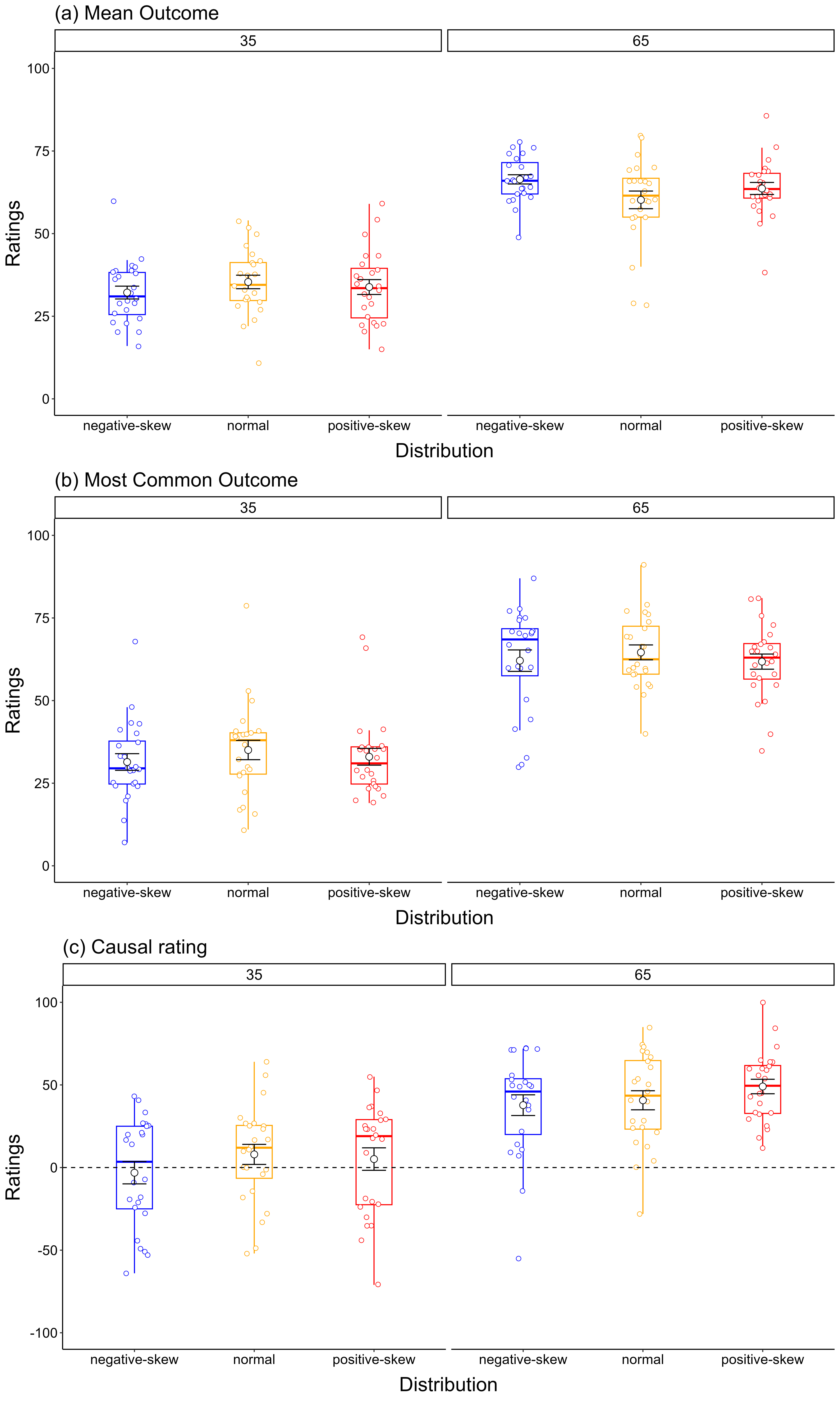

**Experiment 4**

In this experiment, we compared normally distributed outcomes with high (Normal-wide distribution) or low variance (Normal-narrow distribution) against a bimodal outcome distribution at two levels of outcome mean (35 and 65).

***Comparing predictions to observed outcomes***

Participants’ pattern of predictions were again significantly different to the observed outcomes for all cue types. These results are summarised in Table D, and illustrated in Figure D.

**Table D**

*D statistic and bootstrap p value obtained from a Kolmogorov-Smirnov test of equality comparing prediction ratings to observed outcomes in the last 20 presentations of each cue type in Experiment 4.*

|  | 35 | 65 |
| --- | --- | --- |
| Normal Narrow | *D* = .210, *p* < .001 | *D* = .187, *p* < .001 |
| Normal Wide | *D* = .173, *p* < .001 | *D* = .117, *p <* .001 |
| Bimodal | *D* = .150, *p* < .001 | *D* = .178, *p <* .001 |

**Figure D**

*Frequency polygon illustrating the pattern of predictions (histogram) and observed outcomes (solid line) made for different cue types in the final 20 presentations of Experiment 4 average across all participants and separated by cue type.*

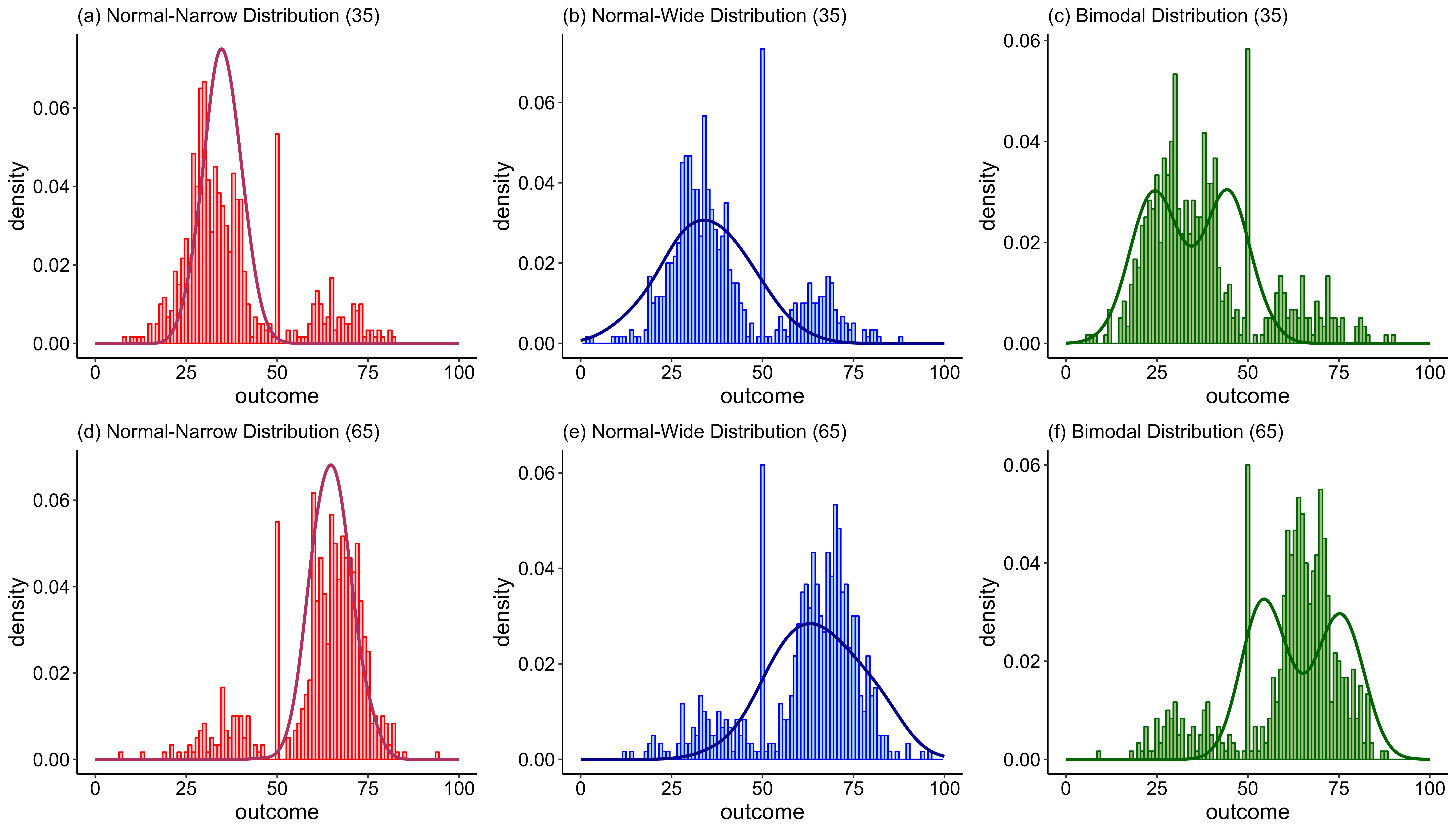

***Test Ratings***

Average outcome estimates in Experiment 4 showed very similar results to that in Experiment 3 (see Figure E). Planned contrasts comparing 1) bimodal outcome distribution to the average of the two normal distributions, 2) normal-narrow vs normal-wide distribution, 3) outcome mean of 35 to outcome mean of 65, and all their interactions, revealed a significant main effect of outcome mean, *F*(1,29) = 71.1, *p* < .001, η_p_^2^ = .710. In this experiment, there was also an overall difference in mean ratings when comparing outcomes sampled from a bimodal vs normal distribution (average over normal-narrow and normal-wide), *F*(1, 29) = 7.31, *p* = .011, η_p_^2^ = .201, suggesting sensitivity to the qualitatively different distribution shapes. No other comparisons reached statistical significance, *Fs*(1,29) $\leq$ 2.59, *p*s $\geq$ .118, η_p_^2^ $\leq$ .082.

Ratings on the most frequent outcome (mode) for each target cue type again showed a sensitivity to the mean, *F*(1,29) = 86.5, *p* < .001, η_p_^2^ = .749, with all ratings not differing significantly from the average mean value of 35 and 65 respectively. Importantly, we again found a significant difference in estimates when comparing bimodal against the two normal distributions, *F*(1, 29) = 3.70, *p* = .064, η_p_^2^ = .113, suggesting some sensitivity to the overall shape of the distribution. No other contrast comparisons or interactions reached statistical significance, *Fs*(1,29) $\leq$ 1.07, *p*s $\geq$ .309, η_p_^2^ $\leq$ .036.

Finally, causal judgements for each of the target treatment cues showed a significant main effect of cue mean, *F*(1,29) = 45.7, *p* < .001, η_p_^2^ = .612, and a marginal difference in ratings for bimodal vs average of the two normal distributions, *F*(1,29) = 3.98, *p* = .05, η_p_^2^ = .121. No other comparisons approached significance, *Fs*(1,29) $\leq$ 2.92, *p*s $\geq$ .098, η_p_^2^ $\leq$ .091. Thus, like in Experiment 3, participants’ causal ratings showed sensitivity to the outcome mean with higher mean producing strong causal beliefs about the efficacy of the treatment. In addition, we found some evidence that mean and mode estimates were also affected by the shape of the distribution outcome values were sampled from, with differences in overall ratings when comparing cues with outcomes sampled from bimodal vs normal distributions, where the differences in distribution may be more pronounced.

### Figure E

*Average ratings (mean denoted by white circle ± standard error) when participants are asked to provide a rating for (a) outcome mean, (b) outcome mode, and (c) causal ratings as a function of actual outcome mean (35 vs 65) and distribution (bimodal, normal distribution with small variance [normal-narrow] and normal distribution with large variance [normal-wide]). Each coloured circle represents an individual participant data point plotted over boxplots illustrating the inter-quartile range and median for each condition.*

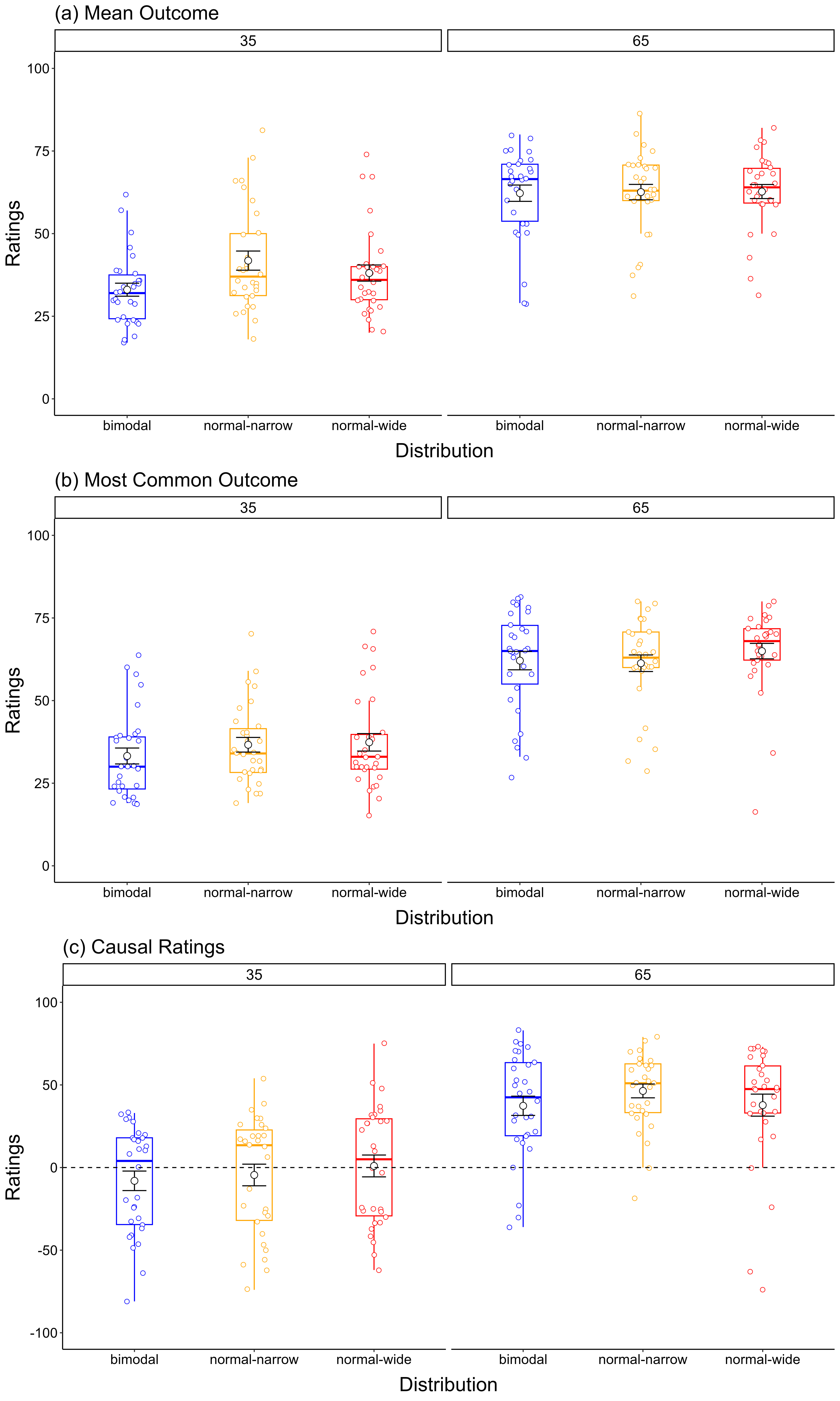

**Computational Modelling Results**

**Model recovery**

To assess the extent to which data generated from each model is distinguishable from the other, we conduct a model recovery analysis. We first generated data using each model, combined with the trial sequences of randomly chosen participants from each experiment, along with their optimal parameter values obtained through model fitting. For each set of simulated participant data, we then ran model fitting with each model with 50 randomly chosen starting points for the parameters. Note that we chose a fully data-informed recovery approach because the Simple Delta and Distributed models are not nested models (they each have additional complexities) and have the same number free parameters (though granted, the Distributed model could be argued to have some inherent complexity that is lacking in the Simple Delta model).

We created 1000 simulated participants per experiment using the Delta Model and a further 1000 using the Distributed Model, each time randomly choosing the trial sequence and optimal parameters values from an actual participant. We then fit both the Delta and Distributed models to each simulated participant’s predictions and chose the best fitting model based on BIC for each simulated participant. Model recovery success was gauged by looking at the proportion of Delta-Model-generated participants that were best fit by the Delta Model, and the proportion of Distributed-Model-generated participants that were best fit with the Distributed model. A breakdown of model recovery results are presented in Table E.

**Table E**

*Percentage of simulated participants recovered by each model, separated by Generating Model and Experiment.*

| Generating Model | Delta | | Distributed | |
| --- | --- | --- | --- | --- |
| Recovered Model | Distributed | Delta | Distributed | Delta |
| Experiment 1 | 78.8 | 21.2 | 95.7 | 4.4 |
| Experiment 2 | 66.3 | 33.7 | 82.9 | 17.1 |
| Experiment 3 | 13.5 | 86.5 | 86.0 | 14.0 |
| Experiment 4 | 5.5 | 94.5 | 80.0 | 20.0 |

*Note: Lightly shaded cells denote cases where the generating model was correctly recovered.*

Overall model recovery success is illustrated in Figure F. Across all experiments, 72.56% of simulated participants were best fit by the model that was used to produce the data. There was a tendency for the distributed model to produce the best fit; 86.15% of Distributed-Model-generated participants were best fit by the Distributed Model whereas only 58.98% of Delta-Model-generated participants were best fit by the Delta Model, and this pattern was particularly distinct in Experiments 1 and 2, where successful recovery of the Simple Delta Model was below 50%.

We interpret this pattern of results to indicate that there are combinations of parameters, perhaps drawn out by the designs of Experiments 1 and 2 in particular (i.e., varying the frequency of different outcome values and the overall base rate through non-target cues rather than manipulating outcome distribution directly), for which the Distributed Model produces data that are not substantially different from what the Delta Model produces. Such manipulations may produce relatively subtle differences in the patterns of predictions generated by the two models. By contrast, Experiments 3 and 4 varied the shape of the outcome distribution itself, contrasting normal against bimodal and skewed distributions. It is therefore unsurprising that the designs of Experiments 3 and 4 proved far more diagnostic, and model recovery for both the Simple Delta and Distributional Model improved substantially. It is also noteworthy that the Distributed Model is capable of generating patterns of data that are not well accounted for by the Delta Model, and also capable of accounting for some patterns in human data which the Delta model cannot.

**Figure F**

*Percentage of Simulated Data Recovered by the Simple Delta and Distributed Model as a function of Generating Model.*

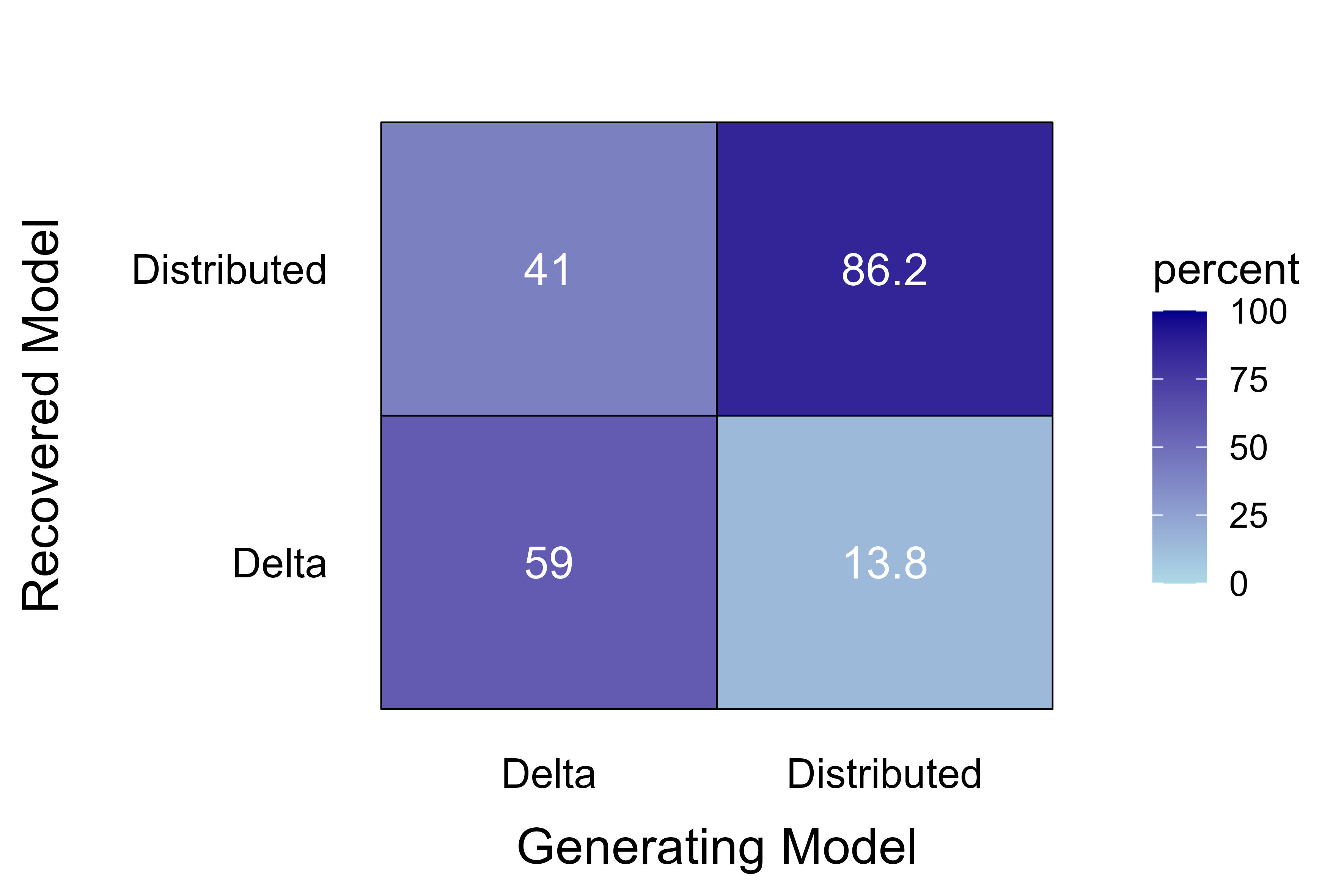
**Parameter Recovery**

We conducted a parameter recovery analysis to determine if the model can recover the true parameters used in data simulation. This analysis allows us to validate whether the fitting procedure can reliably capture the underlying generative process. For each of the 1000 simulations generated by the Delta and Distributed Model for model recovery, we selected the best-fitting parameters of the correct model based on the lowest BIC values obtained across the 50 model fits. To assess parameter recovery, we computed correlations between the true (generating) parameters and the recovered (best-fitting) parameters. Because the 1000 simulations were generated by repeatedly sampling from only each unique participants' trial sequences, many simulations shared identical generating parameters and trial sequences. To account for this structure, we implemented an iterative sampling procedure.

In each iteration, we randomly sampled one simulation per unique participant, ensuring that each participant contributed only once to the correlation analysis for each model. We then computed Pearson correlations between true and recovered parameters separately for each parameter (cue α, context α_x_, softmax temperature *k*, and the model-specific fourth parameter: prediction noise for the Delta model, σ*_e_*, and node activation for the Distributed model, σ*_n_*). This process was repeated 100 times to obtain a distribution of correlation coefficients. We report the median correlation and 95% confidence intervals across these 100 iterations in Experiments 1-4, providing a robust estimate of parameter recovery that accounts for sampling variability (Table F). An example scatterplot illustrating the relationship between true and recovered parameter values in one iteration (iteration number 37, Experiment 4) is presented in Figure G.

**Table F**

*Median Correlation Coefficients with 95% Confidence Interval comparing True (Generating) and Recovered Parameter Values for all Parameters across Four Experiments.*

|  | Experiment 1 | Experiment 2 | Experiment 3 | Experiment 4 |
| --- | --- | --- | --- | --- |
| **Simple Delta**  **Model** | | | | |
| Cue α | 0.24  [-0.14,0.61] | 0.34  [0.19,0.49] | 0.42  [-0.09,0.70] | 0.56  [-0.01,0.83] |
| Context α_x_ | -0.003  [-0.18,0.70] | 0.27  [0.004,0.44] | -0.09  [-0.13,0.80] | -0.04  [-0.07,0.29] |
| Softmax Temp *k* | 0.63  [0.01,0.92] | 0.41  [0.28,0.55] | 0.84  [0.44,0.95] | 0.83  [0.65,0.92] |
| Prediction Noise σ*_e_* | 0.63  [-0.06,0.99] | 0.42  [0.30,0.52] | 0.63  [-0.12,0.86] | 0.32  [-0.22,0.73] |
| **Distributed**  **Model** | | | | |
| Cue α | 0.25  [0.15,0.33] | 0.22  [0.02,0.40] | -0.08  [-0.32,0.22] | 0.10  [-0.22,0.41] |
| Context α_x_ | 0.15  [-0.14,0.67] | 0.67  [0.51,0.84] | 0.40  [-0.12,0.73] | 0.22  [-0.15,0.63] |
| Softmax Temp *k* | -0.01  [-0.33,0.48] | 0.38  [0.22,0.53] | 0.34  [-0.10,0.68] | 0.22  [-0.19,0.54] |
| Node Activation σ*_n_* | 0.80  [0.17,0.95] | 0.84  [0.77,0.90] | 0.75  [-0.08,0.89] | 0.75  [0.04,0.90] |

**Figure G**

*Scatterplot of True vs Recovered Parameter Values obtained from the Delta and Distributed Model in a single iterative sample in Experiment 4. Each point represents a unique participant’s value. Solid line represents the regression line with shaded region depicting SEM.*

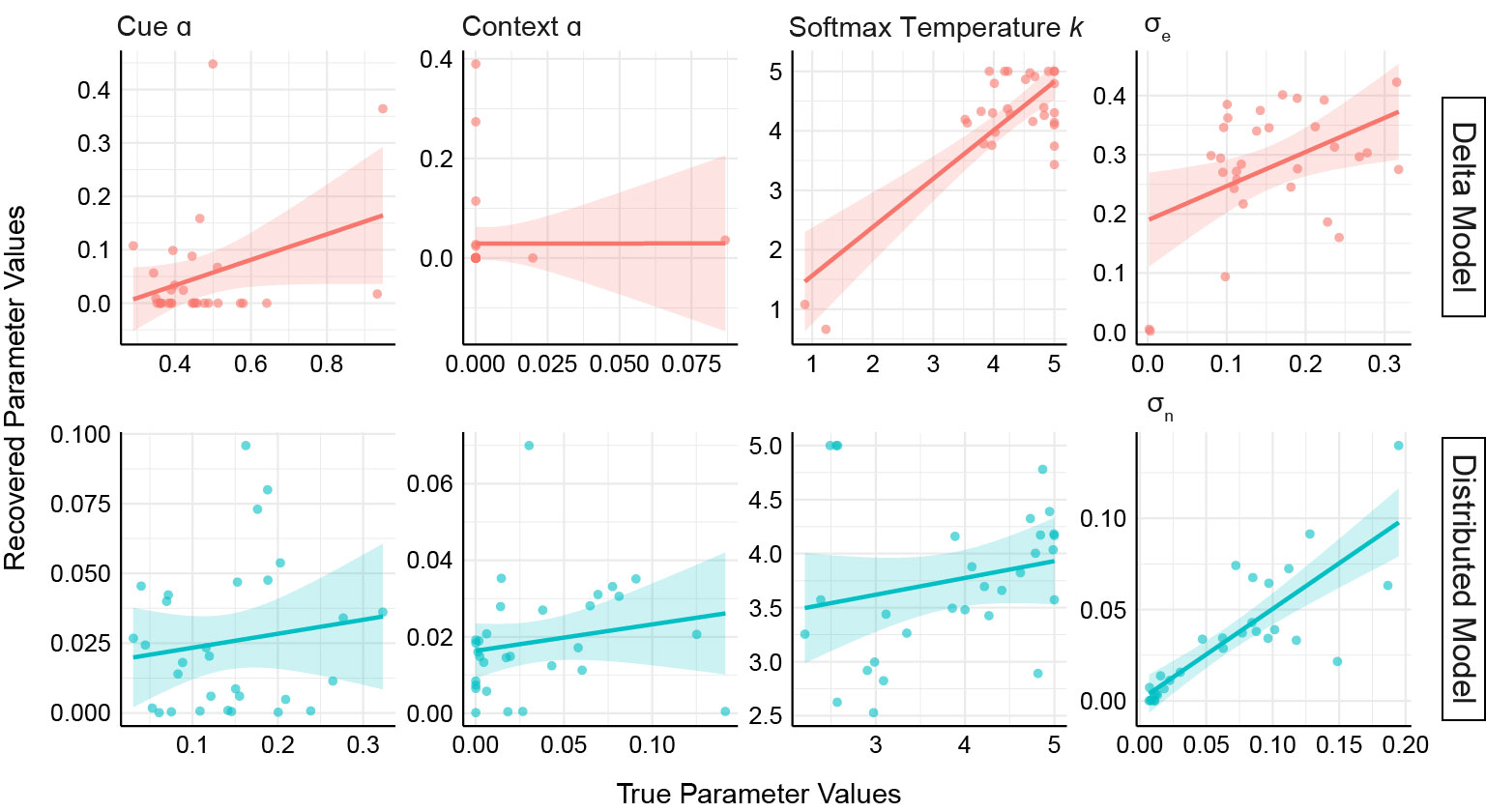

Parameter recovery revealed considerable variability across experiments and parameters. Recovery quality was generally moderate to weak, with some parameters showing wide 95% confidence intervals that included zero or negative values. For the Simple Delta Model, softmax temperature parameter *k* showed the most consistent recovery across experiments (median r = 0.41–0.84), while cue α and context α_x_ showed weakest recovery. For the Distributed Model, σ*_n_* demonstrated the strongest and most consistent recovery across all experiments (median r = 0.75–0.84).

Weak recovery of cue learning rate, α and context learning rate, α_x_ can be attributed to two factors. First, a substantial proportion of recovered parameters fell outside the range of true generative parameter values—only 20.7% of recovered α values fell within the true parameter range for the Delta Model, and 29.7% for the Distributed Model. As noted in Wilson and Collins (2019), parameter recovery can fail when the behavioural data do not provide sufficient constraints to uniquely identify parameter values within the relevant range, even when those parameters were used to generate the data. In generating simulated data for recovery, we defined the starting values of the parameters to match the range of values obtained during empirical model fit, however this restriction did not limit the optimiser’s ability to search for parameter values outside the range –upper and lower boundary values were wide and identical to empirical model fit. Second, for context learning rate α_x_, recovery was weak (median r = -0.09 to 0.27 for Delta; 0.15 to 0.67 for Distributed) because this parameter governs learning about a constant background stimulus that is present on every trial and does not discriminate between cue types. Consequently, α_x_ may produce relatively little trial-to-trial variation in predictions that could distinguish it from other parameters, making it difficult to uniquely identify from behavioural data alone.

Parameters with weak identifiability should be interpreted with appropriate caution when examining individual differences or group-level parameter estimates. The moderate-to-strong recovery of certain parameters (e.g., σ*_n_* in the Distributed model), provides confidence that these specific parameters reflect meaningful individual differences, while parameters with poor recovery should be treated as less informative about the psychological processes they represent. The strong recovery of σ*_n_* is notable as it is unique to the Distributed Model and thus provides important validation for our novel modelling approach. This parameter represents the core theoretical innovation of the Distributed Model, capturing the variability in node activation that produces distributed representations of outcomes. The fact that this parameter shows consistently strong identifiability across all four experiments indicates that the behavioural data contain sufficient information to distinguish distributed from non-distributed learning mechanisms. This suggests that the Distributed Model's architectural features make empirically testable predictions that are reflected in participants' response patterns, even when other shared parameters are less well identified.

More broadly, these results indicate that several model parameters are weakly identified by the behavioural data, meaning that multiple parameter combinations can produce similar patterns of behaviour. This is a common challenge in computational modelling of learning, particularly for: (1) learning rate parameters in tasks with relatively few trials per cue (e.g., 21 intermixed presentations in Experiments 1 & 2), where individual differences in learning rates may be difficult to distinguish from noise; (2) softmax temperature *k*, which may trade off with the model-specific noise parameters (σ_e_ in the Delta model, σ_n_ in the Distributed model) in determining response variability; and (3) learning rate of the context α_x_, attributed to learning about a constant background cue is updated on every trial and does not distinguish between different cue types. Importantly, weak parameter recovery does not necessarily invalidate model comparison or qualitative conclusions about model fit. Poor recovery of individual parameters can occur even when models make clearly distinguishable predictions—for instance, if multiple parameter combinations within a model produce similar behaviour, but the pattern of that behaviour still differs systematically between models. Indeed, our model recovery analysis (reported above) demonstrates that the models themselves can be reliably distinguished in many cases despite weak recovery of individual parameters, suggesting that the key architectural differences between the Simple Delta and Distributed Model learning mechanisms are captured by the data even when precise parameter values are not.

**Analysis of Matching Strategy**

To explore the possibility that participants’ pattern of predictions for the different cues were a result of a direct matching strategy –observed outcomes on trial n are reproduced in participants’ prediction on trial n+1—we ran a simple model comparison for a matching model compared to a non-matching model. The non-matching model assumes simple learning of the outcome mean up to trial n and uses this as a predictor of subsequent prediction ratings.

Both models are defined by a simple linear regression with prediction on trial n+1 as the dependent measure. The matching model has outcome on trial n as the predictor, whereas the non-matching model has the running outcome average up to trial n as a predictor. We then computed a log likelihood ratio of the matching model relative to the non-matching model, and converted the ratios to Bayes Factors in favour of the matching model. A summary of the group-level estimates is shown in Table G. In addition, we ran participant-level analysis by directly correlating predictions on trial n+1 to the observed outcome on trial n. Results from the correlation analysis for each participant can be found on the Open Science Framework. The correlation matrix for participants in Experiment 3 is shown in Figure H.

**Table G**

*Results from model comparison of participants’ predictions based on a matching vs non-matching (learning) model at each level of outcome mean and distribution across all four experiments.*

| Experiment | Mean | Distribution | Likelihood ratio | Bayes Factor (Matching/Non-Matching) |
| --- | --- | --- | --- | --- |
| 1 | 25 | Fixed | NA | NA |
|  | 50 |  | NA | NA |
|  | 75 |  | NA | NA |
|  | 25 | Uniform | 0.9516 | 2.587 |
|  | 50 |  | -2.954 | ^#^.05212 |
|  | 75 |  | 0.5568 | 1.745 |
|  | 25 | Normal | -3.243 | ^#^0.0391 |
|  | 50 |  | -0.168 | 0.8456 |
|  | 75 |  | -10.04 | ^#^.000045 |
| 2 | 15 | Normal | -0.000951 | 0.999 |
|  | 30 | Normal | .2816 | 1.325 |
|  | 50 | Normal | -3.029 | ^#^.0484 |
|  | 70 | Normal | 0.1264 | 1.135 |
|  | 85 | Normal | -4.1661 | 0.155 |
| 3 | 65 | Negative skew | -0.249 | 0.779 |
|  |  | Positive skew | 7.352 | *1558.7 |
|  |  | Normal | 2.491 | *12.07 |
|  | 35 | Negative skew | 4.543 | *93.96 |
|  |  | Positive skew | -8.716 | ^#^.000164 |
|  |  | Normal | -3.978 | ^#^0.0187 |
| 4 | 65 | Normal-Narrow | 0.8552 | 2.352 |
|  |  | Normal-Wide | 1.4629 | *4.319 |
|  |  | Bimodal | -5.5193 | ^#^0.004 |
|  | 35 | Normal-Narrow | 1.9127 | *6.771 |
|  |  | Normal-Wide | 1.4267 | *4.165 |
|  |  | Bimodal | -5.1761 | ^#^0.0057 |

*BF10 > 3, evidence in favour of the matching model

^#^BF01 > 3, evidence in favour of the non-matching model

**Figure H**

*Correlation matrix illustrating the strength of correlation between predictions generated by the Simple Delta (top) and Distributed Model (bottom) on trial n+1 and the observed outcome on trial n for each cue type in Experiment 3. Each column represents data from an individual participant, and each row consists of one of the target cues presented in the study; row labels indicate the distribution type with the outcome mean in parentheses. Cells that appear more red in colour indicate increasingly positive correlation values, whereas cells appearing more blue in colour indicate increasingly negative correlation values, with colours close to white indicating little to no correlation. P-values for statistically significant correlations are included in the relevant cells, whereas correlations which did not reach statistical significance are not shown for visual clarity.*

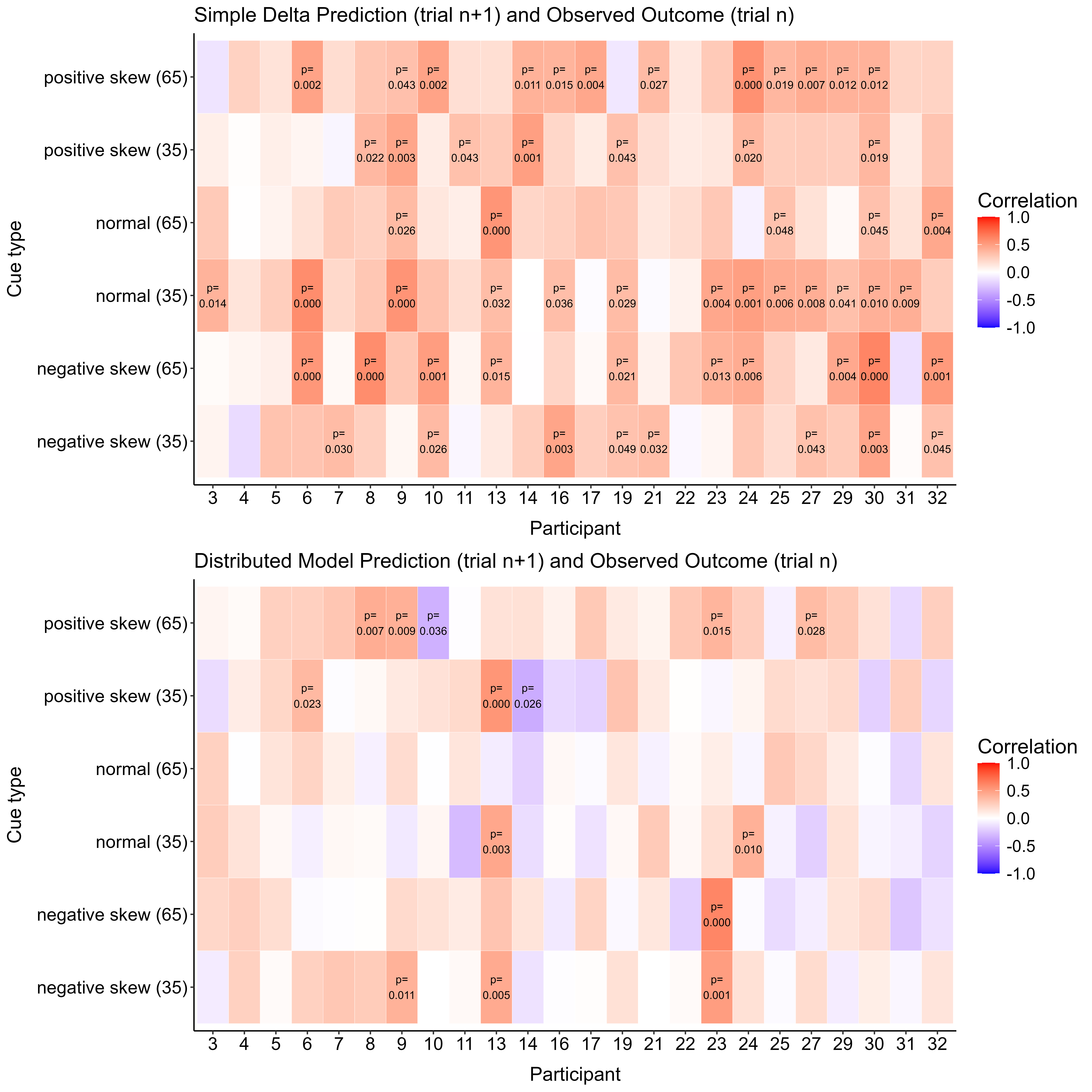
